# DNA repair enzyme NEIL3 enables a stable neural representation of space by shaping transcription in hippocampal neurons

**DOI:** 10.1101/2021.02.09.430416

**Authors:** Nicolas Kunath, Anna Maria Bugaj, Pegah Bigonah, Marion Silvana Fernandez-Berrocal, Magnar Bjørås, Jing Ye

**Affiliations:** Department of Clinical and Molecular Medicine (IKOM), Norwegian University of Science and Technology (NTNU), N-7034, Trondheim, Norway; Department of Microbiology, Oslo University Hospital and University of Oslo, N-0424 Oslo, Norway

## Abstract

DNA repair enzymes are essential for the maintenance of neuronal genome and thereby proper brain functions. NEIL3 is a member of the NEIL family DNA glycosylases initiating oxidative DNA base excision repair. Recent studies show that NEIL3-deficiency leads to impaired spatial performance in mice, decreased adult neurogenesis and altered synaptic composition in the hippocampus. However, it remains elusive how NEIL3 contributes to spatial information coding in hippocampal neurons. Here, we revealed impaired spatial stability in *Neil3^−/−^* CA1 place cells, demonstrating a functional interference of NEIL3 with spatial representations. We identified NEIL3-dependent transcriptional changes in response to spatial exploration and defined its regulatory role specifically for NMDA receptor subunits and immediate early genes. Our work demonstrates a non-canonical role of NEIL3 in modulating the functional plasticity of place cells by shaping the neuronal transcriptome, thus sheds light on the molecular determinants enabling a stable neural representation of space.

## INTRODUCTION

Due to the high oxidative load and free radicals produced by cellular metabolism in the brain, repair of oxidative DNA damages in neurons is extremely important for the maintenance of proper brain functions (*1*). The NEIL3 DNA glycosylase is one of the essential enzymes that initiates the Base Excision Repair (BER) by recognizing and removing oxidative DNA base lesions (*2–4*). NEIL3 displays discrete expression patterns in the brain and is enriched in neurogenic niches (*4, 5*). Recent studies demonstrate the essential role of NEIL3 in proliferating neurons in newborn mice as well as continuous adult neurogenesis in the hippocampus (*6, 7*). Mice lacking NEIL3 have a normal lifespan without a predisposition to cancer or increased spontaneous mutation frequencies (*8*). However, *Neil3^−/−^* mice display a differential synaptic composition in the hippocampus and an impaired spatial learning phenotype in the Morris Water Maze (*6*), suggesting a distinct role of NEIL3 in the regulation of hippocampal functions.

The hippocampus plays a pivotal role in encoding and recall of spatial and nonspatial episodic memories. A high degree of transcriptional variety has been delineated in subregions and sub-cell-populations of the hippocampus (*9–13*). Cell type-specific and activity-induced changes in gene transcription have also been described to promote the refinement and plasticity of neural circuits for cognition and behavior (*14*). Experience-dependent transcriptome has been characterized in hippocampal dentate granular cells (*15*) and the comprehensive transcriptional and epigenomic landscapes have been mapped during memory formation and recall in hippocampal engram ensembles (*16*). The study of hippocampal function has been revolutionized since the discovery of place cells, one of the most remarkable neuronal correlates for spatial cognition and behavior (*17, 18*). Place cells are neurons that encode spatial information in their firing patterns (place maps), identified in all sub-regions of the hippocampus (*19–21*). Over the decades, the activity of place cells has been extensively studied in response to spatial experience, sensory input and task demands (*22*). However, it is far less understood how epigenomic and transcriptomic interplay regulates the functional properties of hippocampal neurons.

In this study, we elucidated a novel function of NEIL3 in hippocampal CA1 pyramidal cells at both the functional and molecular levels. We studied functional properties of CA1 neurons in *Neil3^−/−^* mice during spatial exploration, discovering impaired spatial stability in CA1 place cells lacking NEIL3. We identified spatial experience-induced gene expression in *Neil3^−/−^* CA1 neurons, highlighting NEIL3-dependent gene modulation in epigenetic and synaptic regulation. We defined experience-dependent regulation of NMDA receptor subunits and immediate early genes, implicating a role of NEIL3 in the molecular correlates of constrained spatial memory. Our work provides evidence that NEIL3-dependent transcriptional interplay is involved in shaping the functional plasticity of place cells.

## RESULTS

### *Neil3^−/−^* CA1 place cells displayed normal spatial activity

Place cells encode spatial information in their environment-specific firing patterns (“place fields”) (*19*). To explore whether NEIL3 contributes to the spatial information coding in hippocampal neurons, we recorded CA1 place cell activity in wildtype and *Neil3^−/−^* mice while the animals were freely moving in an open field environment (Fig. 1A). The implant locations were evenly distributed along the proximodistal axis of CA1 in both genotypes (Fig. 1C), as space is represented non-uniformly along the transverse axis of CA1 (*23*). Place cells were defined as cells with scores for spatial information content passing the 95^th^ percentile of a distribution for randomly shuffled data from all recorded CA1 cells within the group (Fig. 1D, spatial information content above 0.554 for wildtype and above 0.547 for *Neil3^−/−^*). Based on this criterion, we identified a total of 355 place cells (85% of 419 putative principal neurons recorded) in wildtype (n=4) and 313 place cells (78% of 402 putative principal neurons recorded) in *Neil3^−/−^* (n=4) mice. Most CA1 place cells had a single environment-specific firing pattern (the “place field”), but more than one place field was also observed in some cases (Fig. 1B and Fig. S1). A higher fraction of place cells in *Neil3^−/−^* CA1 displayed multiple firing fields (29% vs 18% in wildtype, Fig. 1F). The average number of place fields differed significantly (wt: 1.22±0.03, *Neil3^−/−^*: 1.39±0.04 [mean±SEM], p=0.0001 unpaired t-test with Welch’s correction and pcorr. =0.0441 nested t-test). We also assessed a range of electrophysiological characteristics for all place cells in the wildtype and *Neil3^−/−^* groups (Figure 6E). No difference was observed regarding spatial information content (wt: 1.19±0.02, *Neil3^−/−^*: 1.16±0.03, mean±SEM across cells), spatial coherence (wt: 0.88±0.02, *Neil3^−/−^*: 0.88±0.02) or within-session spatial stability (wt: 0.75±0.01, *Neil3^−/−^*: 0.71±0.01). Minor differences were observed in the mean firing rate (wt: 0.96±0.04Hz, *Neil3^−/−^*: 0.79±0.04Hz, p=0.0012 and pcorr.=0.3645), peak firing rate (wt: 6.31±0.23Hz, *Neil3^−/−^*: 5.61±0.23Hz, p=0.0375 and pcorr.=0.5519) and mean field size (wt: 458±13mm^2^, *Neil3^−/−^*: 412±014mm^2^, p=0.0181 and pcorr.=0.3245) (Fig. 1E). These observations demonstrate generally normal spatial activity of CA1 place cells in mice lacking NEIL3.

**Figure 1.**
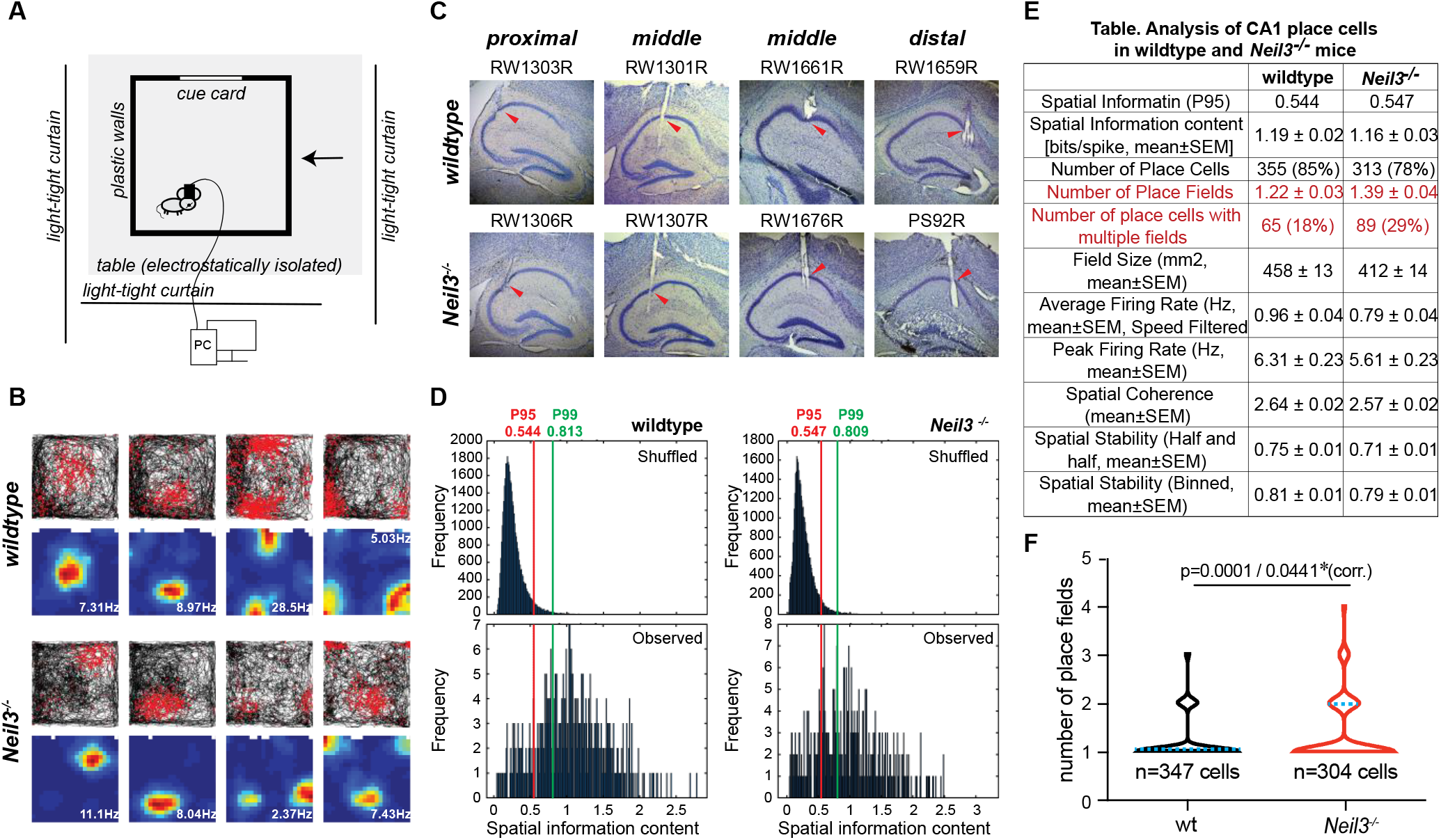
CA1 place cells recorded in wildtype and *Neil3^−/−^* mice. (**A**) Schematic illustration of the experimental set-up for extracellular neuronal live recordings. (**B**) Firing pattern of representative place cells recorded in wildtype and *Neil3^−/−^* CA1. Trajectory maps with superimposed spike locations (red dots) are shown at top and color-coded rate maps (red for peak and blue for firing rates) with indicated peak firing rate are shown at bottom. (**C**) Cresyl-violet stained hippocampal sections showing the tetrode locations in all implanted animals (n=4 wildtype and n=4 *Neil3^−/−^* mice). (**D**) Distributions of observed spatial information scores and the randomly shuffled data for the entire cell samples recorded in wildtype or *Neil3^−/−^* CA1. The 95^th^ and the 99^th^ percentile of the shuffled distribution are marked by the red and green line. The 95th percentile was used as a threshold to define place cells. (**E**) Table of general electrophysiological properties of place cells in wildtype versus *Neil3^−/−^* mice. Significant differences were observed in *Neil3^−/−^* CA1 regarding the number of place fields per cell and the fraction of place cells with multiple place fields, marked in red. (**F**) Violin plot illustrating the number of place fields per place cell in wildtype and *Neil3^−/−^* CA1(blue line indicates median). The average number of place fields differed significantly between genotypes (pcorr.=0.0441, nested t-test).

### *Neil3^−/−^* CA1 place cells displayed impaired spatial stability

Hippocampal place cells are able to maintain a stable spatial map in the familiar environment and alter their firing patterns upon environmental changes (termed “remapping”) (*24, 25*). When animals are exposed to a novel environment, place fields of specific cells may change in firing rate, shift in location, appear or disappear, a process known as “global remapping” (*26*). We recorded CA1 place cells in wildtype and *Neil3^−/−^* mice over five sequential sessions in the familiar or novel environments consisting of a black or white colored square recording chamber (see methods and Fig. 2A). Spatial global remapping was measured by cross-correlation of rate maps from the same cell recorded in two different environments (familiar vs novel). Both wildtype and *Neil3^−/−^* place cells reliably remapped to the new environment with similar low correlation coefficients in the whole population (A1 vs B1, wt - 0.028±0.026 vs *Neil3^−/−^* 0.030±0.024 [mean±SEM], p>0.1, Fig. 2A-B and Fig. S2B), suggesting that NEIL3-dependent transcriptional and thereafter synaptic changes are not essential for the remapping of CA1 place cells.

**Figure 2.**
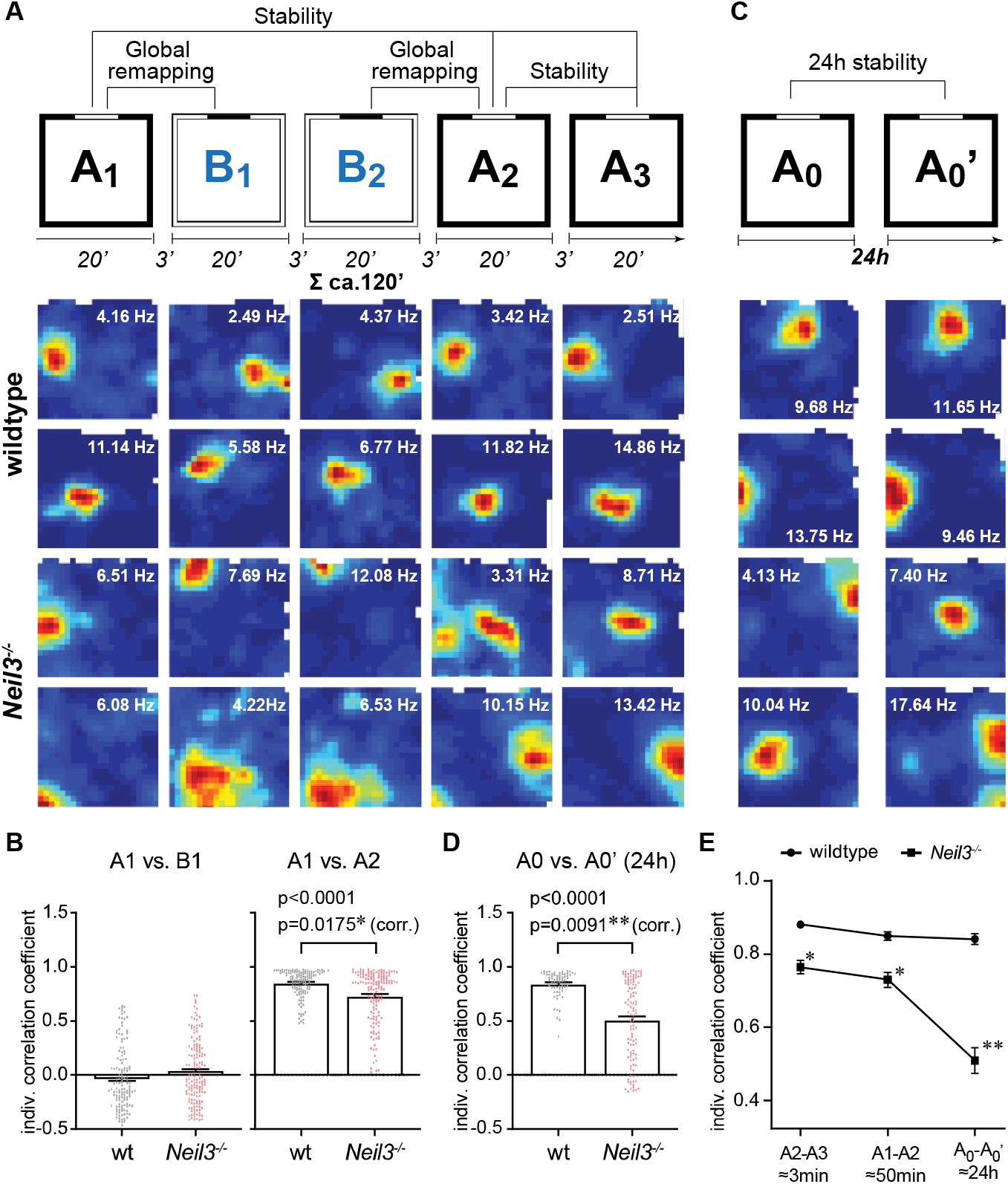
*Neil3−/−* CA1 place cells displayed impaired spatial stability. (**A**) CA1 place cells in wildtype and *Neil3^−/−^* mice were recorded for five sessions (20 min of recording and 3 min of rest) in a sequence of familiar (Room A) and novel (Room B) environments (top row). Global remapping of place cells was observed in both genotypes from the familiar environment A (trial in A1) to the novel environment B (trials in B1 and B2). Wildtype place cells widely retrieved original maps when re-exposed to the familiar environment (trials in A2 and A3), whereas a proportion of *Neil3^−/−^* place cells kept generating new maps. Rate maps of two representative place cells in wildtype and *Neil3^−/−^* CA1 are shown and peak firing rates are indicated. (**B**) Correlation coefficients between trials in A1 and B1 or in A1 and A2 were analyzed for the whole population of wildtype and *Neil3^−/−^* place cells. The firing patterns of place cells in the familiar environment A were re-tested after one day (A0 and A0’, ca. 24-hour interval, as illustrated). Wildtype place cells largely kept the same firing patterns in both trials, whereas *Neil3^−/−^* place cells often generated new maps in the second trial on Day 2. Rate maps of two representative place cells in wildtype and *Neil3^−/−^* CA1 are shown and peak firing rates are indicated. Correlation coefficients between trials in A0 and A0’ were analyzed for the whole population of wildtype and *Neil3^−/−^* place cells that were monitored over the course of 24h. (**E**) The line graph shows the deteriorated spatial correlation of *Neil3^−/−^* place cells in the familiar environments over a longer time course. Statistics were conducted using unpaired t-test with Welch’s correction at the population level (each cell as statistical unit) and using a nested t-test at an animal level (pcorr., each animal as statistical unit, n=4 for each genotype).

Further, we assessed spatial stability by comparing rate maps from the same cell recorded in the same familiar environment A but different sessions. 2.3% of place cells in wildtype, but 18.5% in *Neil3^−/−^* CA1, expressed correlation coefficients lower than 0.5 across two familiar environments (A1 vs A2, 50-min interval). The mean correlation coefficient was significantly lower in the population of *Neil3^−/−^* place cells (wt 0.851±0.011 vs *Neil3^−/^* 0.731±0.020 [mean±SEM], p<0.0001 and pcorr.=0.0175), indicating that a larger fraction of *Neil3^−/−^* place cells had shifted place maps in the familiar environment after two novel environment trials (Fig. 2A-B, Fig. S2C). Lower spatial correlation was also observed in *Neil3^−/−^* place cells recorded in two sequential sessions (A2 vs A3, ca. 3-min interval) with 14.8 % of cells having correlation coefficients below 0.5 (0% of wildtype cells, Fig. S2C, wt 0.882±0.011 vs *Neil3^−/−^* 0.765±0.018 [mean±SEM], p<0.0001 and pcorr.=0.0235). These results suggest impaired spatial stability in CA1 place cells lacking NEIL3.

To further assess the long-term spatial stability of *Neil3^−/−^* place cells, we monitored the firing patterns (place maps) of place cells in the same familiar environment for two days with a 24-hour interval. We reliably recorded 68 wildtype and 113 *Neil3^−/−^* place cells over two days (all cells listed in Fig. S3). A further increased fraction of *Neil3^−/−^* place cells (44.4%) did not retrieve the original place maps in the same environment recorded on day 2 (1.5% of wildtype cells, correlation coefficients below 0.5) with a significantly lower correlation coefficient in the whole population (Fig. 2C-D, wt 0.842 ± 0.015 vs *Neil3^−/−^* 0.509 ± 0.036 [mean±SEM], p<0.0001 and pcorr.=0.009). The spatial correlation of *Neil3^−/−^* place cells in the familiar environments was remarkably deteriorated when cells were recorded over a longer time course (Fig. 2E, Fig. S2D, 0.765±0.018 [A2vsA3, ca.3min], 0.731±0.020 [A1vsA2, ca.50min], 0.509±0.036 [24h], p<0.0001 for both A1A2 vs. A0A0’ and A2A3 vs. A0A0’, mixed model, Sidak’s multiple comparisons test). These results demonstrated that *Neil3^−/−^* place cells displayed impaired long-term spatial stability, linking to the decreased spatial performance observed in *Neil3^−/−^* mice (*6*).

### NEIL3-dependent differentially expressed genes (DEGs) are overrepresented in synaptic components of CA1 neurons and associated with a broad spectrum of neural biological processes

To reveal NEIL3-dependent molecular underpinnings required for the functional plasticity of CA1 neurons, we explored differential gene expression in *Neil3^−/−^* versus wildtype CA1 at baseline (no behavioral intervention, Fig. 3) as well as after a spatial exploration paradigm, representing a similar behavioral setup as used for the place cell recording (Fig. 4). Whole-transcriptome sequencing (RNAseq) was performed using RNA samples from laser-dissected dorsal CA1 (stratum pyramidale) and differential gene expression in *Neil3^−/−^* CA1 cells was analyzed using DESeq2 (*27*).

**Figure 3.**
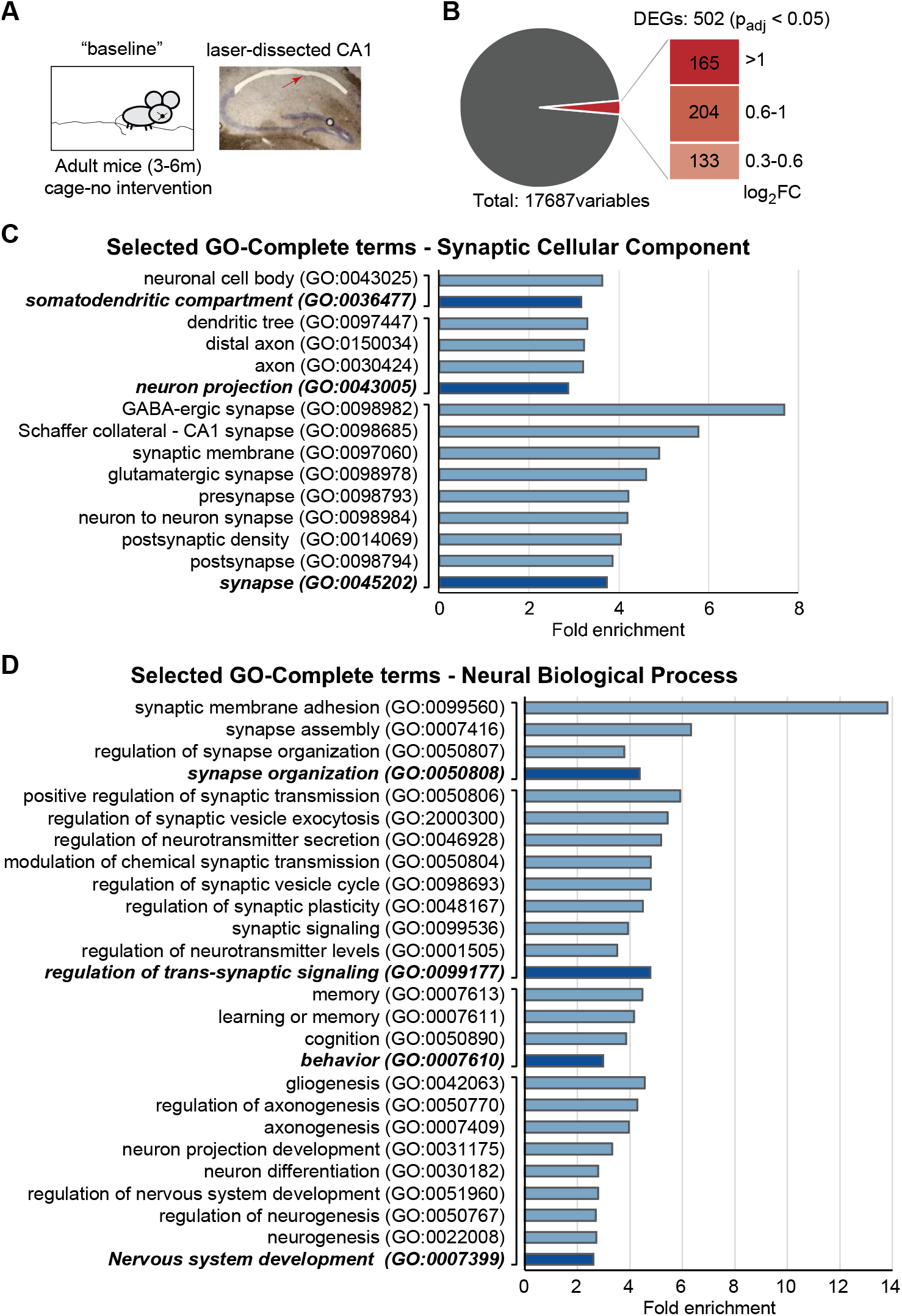
Gene Ontology overrepresentation analysis of differentially expressed genes (DEGs) in *Neil3−/−* CA1 neurons without behavior intervention. (**A**) RNA samples were prepared from laser dissected CA1 of wildtype and *Neil3^−/−^* mice without behavioral intervention (baseline, n=2 per genotype). (**B**) Pie chart showing the proportion of DEGs (Benjamini-Hochberg p_adj_ < 0.05) passing different thresholds of log_2_FC. 502 DEGs passing the log_2_FC ≥ 0.3 were used in PANTHER Gene Ontology (GO) overrepresentation analysis. (**C**) Selected GO-Complete terms of synaptic cellular components overrepresented (Binomial, Bonferroni, FDR < 0.05) in the DEGs. (D) Selected GO-Complete terms of neural biological processes overrepresented (FDR < 0.05) in the DEGs. The dark bars represent the ancestral terms, and the light bars represent the child terms.

**Figure 4.**
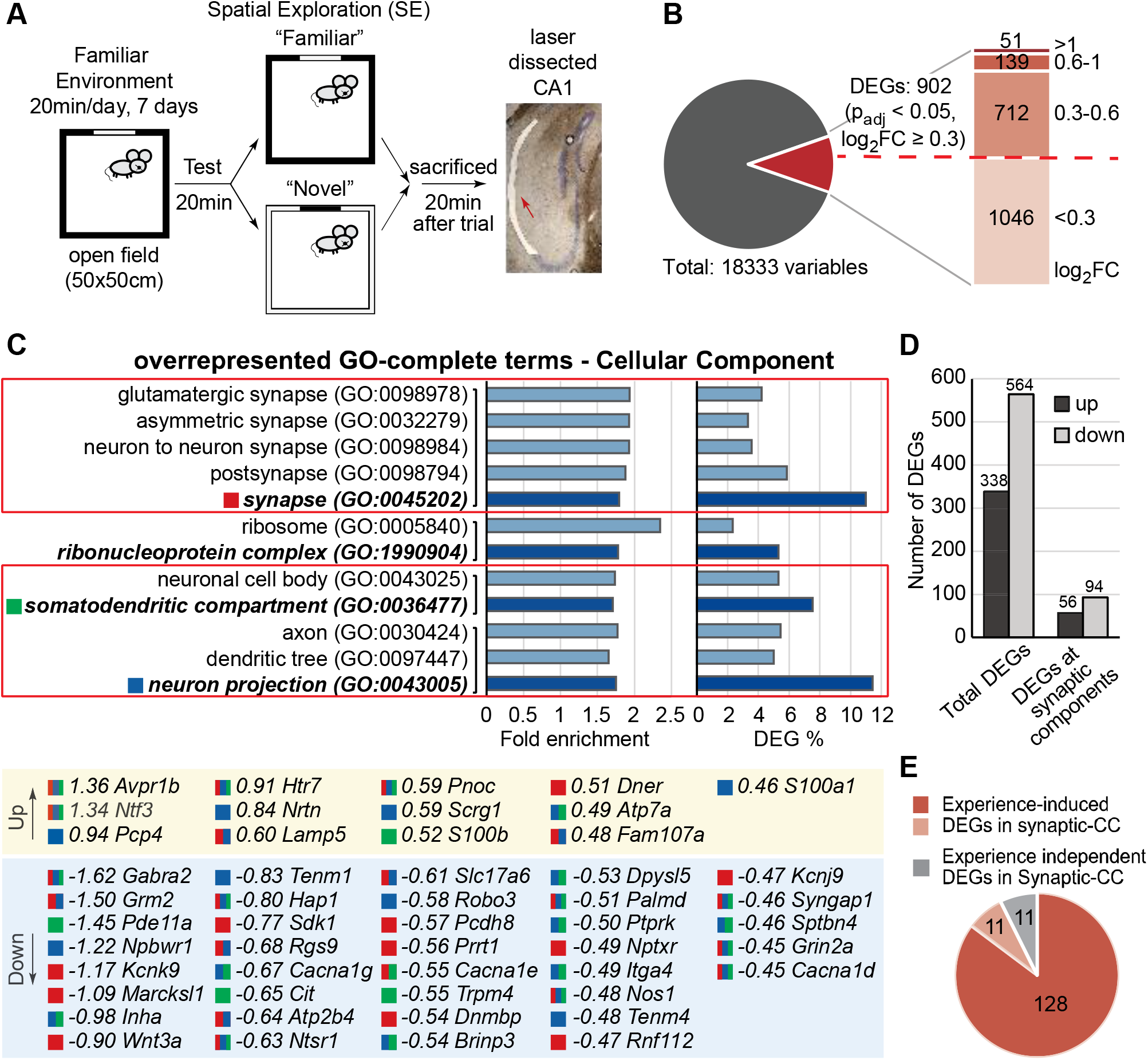
Overrepresentation of NEIL3-dependent differentially expressed genes (DEGs) at synaptic cellular components after spatial exploration. (**A**) Schematic illustration of the experimental design. “Familiar” and “Novel” were combined as one spatial exploration (SE) condition with n=5 for wildtype and n=6 for *Neil3^−/−^*. (**B**) Pie chart showing the proportion of DEGs (Benjamini-Hochberg p_adj_ < 0.05) passing different thresholds of log_2_FC. 902 DEGs passing the log_2_FC ≥ 0.3 (separated by the red-colored dotted line) were used in a Gene Ontology (GO) overrepresentation analysis. (**C**) GO-Complete Cellular Component (CC) terms overrepresented (FDR < 0.05) in the DEGs. The dark bars represent the ancestral terms, and the light bars represent the child terms (full list in supplement table 3). Three ancestral terms (highlighted and color-coded), “synapse”, “neuron projection” and “somatodendritic compartment”, were defined as terms for synaptic components. The top 50 NEIL3-dependent DEGs (log_2_FC values were indicated, marked with up or down-regulation) overrepresented in different synaptic cellular components are listed below. (**D**) Bar-plot showing the proportion of up-and down-regulated DEGs in total and DEGs annotated with synaptic function related GO-CC terms. (**E**) Pie chart showing the proportion of experience dependent DEGs overrepresented in GO-CC terms related to synaptic components, in which 128 DEGs were not differentially regulated in the baseline condition (red), 11 DEGs were inversely regulated (pink) and 11 were similarly regulated (grey) in the baseline condition.

At the baseline condition, a total of 502 genes that passed the criterion of p_adj_<0.05/log_2_FC ≥ 0.3 (fold change ≈ 1.23) were defined as the DEGs in *Neil3^−/−^* CA1 (Fig. 3B, volcano plot in Fig. S4A). We chose a relatively low log_2_FC threshold (0.3) to include all possible NEIL3-dependent genes that are potentially biologically relevant. 60% of DEGs (303 out of 502 DEGs) were upregulated and 40% (199 out of 502 DEGs) were downregulated. Within the body of differentially regulated genes, 165 genes passed the log_2_FC ≥ 1 (fold change 2) and 369 genes passed the log_2_FC ≥ 0.6 (fold change ≈ 1.51). All 502 DEGs were analyzed using the Gene Ontology (GO) consortium/PANTHER classification system (*28*). A statistical overrepresentation analysis of GO terms for the attributed cellular components (CC) and the associated biological processes (BP) as well as PANTHER terms for protein class (PC) was performed (Fig. S4 and supplement table 1-2). We highlighted the respective parent terms that were located higher in the loose GO hierarchy, even though subordinate, more specialized child terms formally had a larger statistical overrepresentation (“fold enrichment”). NEIL3-specific DEGs were overrepresented in GO-CC-terms referring to the parent term “synapse” (GO:0045202, fold enrichment 3.74), including sub-terms such as pre- and post-synapse, GABAergic and glutamatergic synapse, synaptic membrane and postsynaptic density. The parent terms “neuron projection” (GO:0043005, fold enrichment 2.89) and “somatodendritic compartment” (GO:0036477, fold enrichment 3.18) were also significantly enriched, including the sub-terms “axon” (GO:0030424), “dendritic tree” (GO:0097447) as well as “neuronal cell body” (GO:0043025) (Fig. 3C, heat plots of individual genes in Fig. S5). The GO-CC analysis refers to the subcellular location of NEIL3-dependent DEGs overrepresented in synaptic components suggesting a functional relevance of NEIL3 for synaptic regulation. This is supported by a GO-BP analysis, showing a high overrepresentation of NEIL3-dependent DEGs in synaptic processes, such as regulation of trans-synaptic signaling (GO:0099177, fold enrichment 4.78), synapse organization (GO:0050808, fold enrichment 4.37) and their respective sub-terms (Fig. 3D). Nevertheless, NEIL3-dependent DEGs were associated with a broad spectrum of biological processes, related to behavior, development, ion transport and blood circulation (Fig. S4D, supplement table 2).

### NEIL3 impacts spatial experience-induced gene expression associated with synaptic and epigenetic regulation

In the “spatial exploration” condition, wildtype and *Neil3^−/−^* mice were habituated to an open field (“familiar”) environment for 20 min daily in a sequence of 7 days and tested in the familiar or a novel environment before termination (Fig. 4A). This sequence from a familiar to a novel environment was identical to the one used to induce global remapping of CA1 place cells as shown in Fig. 2. However, no differential gene expression was detected in the wildtype or the *Neil3^−/−^* group between these conditions (“familiar” vs “novel”, all genes had a Padj ≈ 1 with DESeq2). We therefore merged samples tested in the familiar and novel environments to one “spatial exploration” (SE) condition, within which we compared both genotypes. In this case, a total of 902 genes passed the criterion of P_adj_<0.05/log_2_FC ≥ 0.3, including 51 genes passed log_2_FC ≥ 1 and 190 genes passed log_2_FC ≥ 0.6, and were defined as NEIL3-dependent DEGs in response to spatial exploration (Fig. 4B, volcano plot in Fig. S6A). Within this body of differentially regulated genes, 37% (338 out of 902 DEGs) were upregulated and 63% (564 out of DEGs) were downregulated.

NEIL3-associated DEGs after spatial exploration were highly overrepresented in GO-CC terms related to the parent terms of “synapse” (GO:0045202, fold enrichment 1.79, 99 out of 902 DEGs), “neuron projection” (GO:0043005, fold enrichment 1.75, 103 out of 902 DEGs), “somatodendritic compartment” (68 out of 902 DEGs, GO:0036477, fold enrichment 1.71) and “ribonucleoprotein complex” (GO:1990904, fold enrichment 1.78, 48 out of 902 DEGs) (Fig. 4C, full list in supplement table 3). A total number of 150 DEGs (56 upregulated and 94 downregulated, 17% of total DEGs) were part of GO-CC terms related to synaptic components (genes enriched in GO-CC-terms “synapse” and/or “neuron projection” and/or “somatodendritic compartment”) (Fig. 4C-D). Within the 150 DEGs at synaptic components, more than 90 % were dependent on the animals’ behavioral experience in the spatial environment (Fig. 4E, 128 genes not differentially regulated and 11 inversely regulated in the baseline condition). Again, this was supported by a GO-BP analysis, showing a high overrepresentation of NEIL3-dependent DEGs in synaptic processes, such as regulation of trans-synaptic signaling (GO:0099177, fold enrichment 2.1), synapse organization (GO:0050808, fold enrichment 2.6) and their respective sub-terms (Fig. 5B, full list in supplement table 4). A total of 62 DEGs were associated with synaptic processes (Fig. 5B, also see heat plots of all 62 genes in Fig. S6B), among which the majority (82%) were not recognized in *Neil3^−/−^* CA1 at baseline. Taken together, these results suggest a functional relevance of NEIL3 for the experience-induced synaptic regulation.

**Figure 5.**
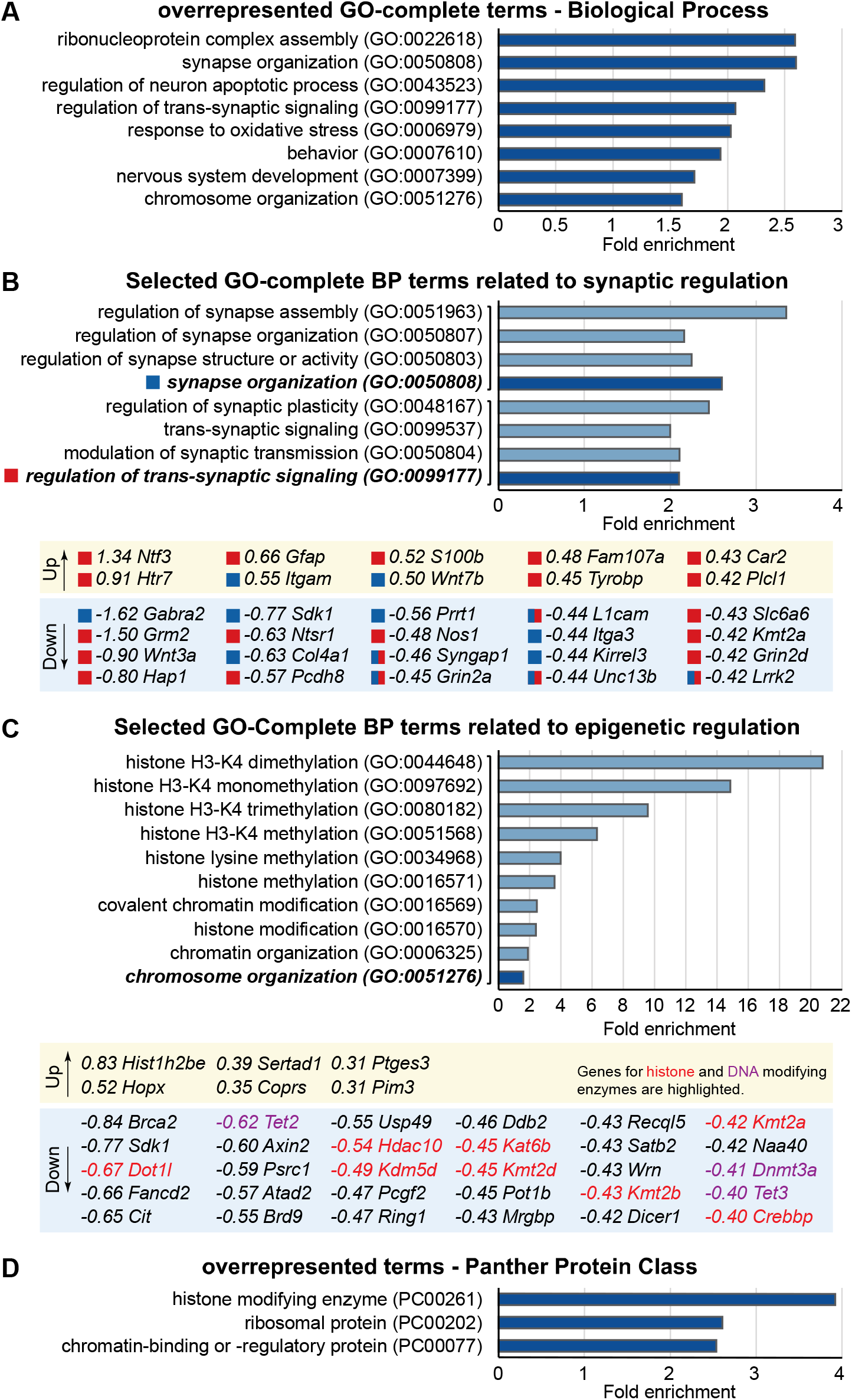
Overrepresentation of NEIL3-dependent DEGs at distinct biological processes after spatial exploration. (**A**) GO-Complete Biological Process (BP) terms overrepresented (FDR < 0.05) in the DEGs. Only ancestral terms are listed (full list in supplement table 4). Selected GO-Complete BP terms related to synaptic regulation. The dark bars represent the ancestral terms, and the light bars represent the child terms. The ancestral terms, “synapse organization” and “regulation of trans-synaptic signaling”, are highlighted and color coded. The top 30 NEIL3-dependent DEGs (log_2_FC values are indicated and marked with up-or down-regulation) overrepresented in respective BP terms are listed below. (**C**) Selected GO-Complete BP terms related to epigenetic regulation. The dark bar represents the ancestral term, and the light bars represent the child terms. All DEGs overrepresented in these BP terms are listed below. Genes for histone (red) and DNA (purple) modifying enzymes are highlighted. (**D**) PANTHER Protein Class **D**terms overrepresented (FDR < 0.05) in the DEGs.

Interestingly, we noted that in response to spatial experience, a number of chromatin-modifying enzymes showed differential expression in NEIL3-deficient mice. We identified DEGs overrepresented in terms of Panther Protein Class “histone modifying enzyme” (PC00261, fold enrichment 3.92) and “chromatin-binding or -regulatory protein” (PC00077, fold enrichment 2.54) (Fig. 5D) as well as in GO-BP parent terms “chromosome organization” (GO:0051276, fold enrichment 1.6) (Fig. 5C). A set of genes (58 out of 902 DEGs) essential for the DNA and histone methylation processes, such as histone lysine methyltransferases (*Kmt2*, *Kmt5*, *Dot1l*), Histone deacetyltransferase (*Hdac10*), DNA methyltransferase (*Dnmt3a*) and DNA methylcytosine dioxygenase (*Tet2* and *Tet3*), were discovered (Fig. 5C, also see heat plots in Figure S6C). This observation suggests that NEIL3 impacts experience-induced epigenetic regulation in hippocampal CA1 neurons.

### NEIL3-deficiency causes changes in regulatory NMDA-receptor subunits

NEIL3 deficiency leads to experience-dependent transcriptional changes of CA1 genes relevant to synaptic regulation with a particular emphasis on glutamatergic signaling. The transcription of NMDA-receptor 2 (NR2) subunits, such as NR2A (*Grin2a*, log_2_FC=−0.45) and NR2D (*Grin2d*, log_2_FC=-0.42), as well as the neuronal nitric oxide synthase (*Nos1*, log_2_FC=-0.48) were significantly downregulated in *Neil3^−/−^* CA1 after spatial exploration (Fig. 5B and S6). NR2A is a regulatory NMDAR subunit that has been implicated in spatial learning and memory-related processes together with the associated subunit NR2B (*29–32*). The neuronal NOS1 has been shown to interact with NMDAR signaling via PSD-95 (*33–35*). We therefore assessed the expression of the regulatory NMDAR subunits (NR2A and NR2B) as well as PSD-95 in *Neil3^−/−^* CA1 neurons by immunohistochemistry (IHC) at baseline and on test day 7 after a 6-day habituation period that largely in parallel to the paradigm used for our RNAseq experiments. The confocal immunofluorescent images showed punctuate fluorophore labeling of NR2A, NR2B and PSD95 in the CA1 pyramidal layer (Fig. 6A-B and Fig. S7A). NR2A, but not NR2B, was significantly upregulated in CA1-SP (stratum pyramidale) in response to spatial experiences (Fig. 6C, p=0.0050 for wt, p=0.0017 for *Neil3^−/−^*). No difference was detected regarding the immunoreactivity of PSD95 after spatial exploration in both genotypes (Fig. S7A-B). Similar spatial exploration induced upregulation of NR2A was also observed in the stratum oriens (SO) and stratum radiatum (SR) of wt and *Neil3^−/−^* CA1 (Fig. S7C-D). Strikingly, we identified an increased ratio of NR2A and NR2B in the CA1 region of NEIL3-deficient mice in response to spatial exploration (Fig. 6C and Fig. S7C-D). An increased NR2A:NR2B ratio has been associated with constrained long-term spatial memory (*36*), which is consistent with previous behavior studies showing an impaired spatial learning and memory performance in NEIL3-deficient mice (*6*).

**Figure 6.**
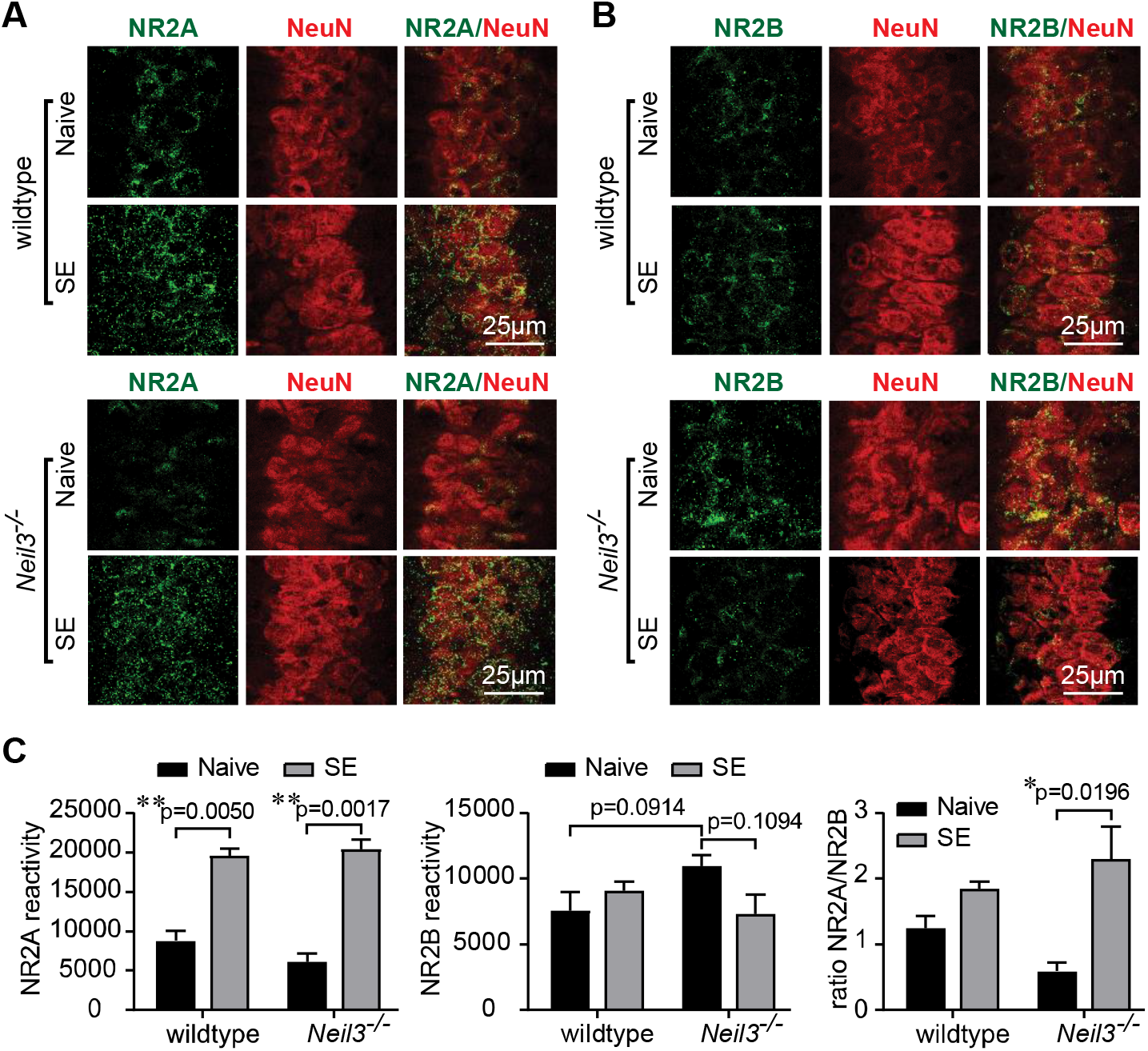
The composition of regulatory NMDA-receptor subunits at CA1 was altered in NEIL3-deficienct mice. Confocal images showing dorsal hippocampal CA1 neurons that were immuno-stained with antibodies against NR2A (green) and NeuN (red) in (**A**) or NR2B (green) and NeuN (red) in (**B**) from wt and *Neil3^−/−^* mice without (Naive) or after **<u>s</u>**patial **<u>e</u>**xploration (“SE”). Bar diagrams showing the immunoreactivity (quantified in Imaris, see methods) of NR2A or NR2B at the pyramidal layer of CA1 and the ratio of NR2A:NR2B was calculated for each animal individually. Statistics were conducted at animal level (3 animals with 2 hippocampal slices for each genotype, two-way-ANOVA, Sidak’s correction, error bars indicate SEM).

### NEIL3 modulates spatial experience induced expression of immediate early genes

Immediate early genes (IEGs) have been identified as key components in synaptic plasticity and as cellular representations in neuronal ensembles underlying the memory trace/engram (*37, 38*). Induced expression of IEGs such as *Arc* and *c-Fos* has been found in hippocampal CA1 neurons associated with neuronal activity during spatial learning (*39–42*). In our transcriptome data, we observed upregulation of several IEGs, including *Arc* and *c-Fos*, following spatial exploration (SE) in the wildtype as well as *Neil3^−/−^* CA1, but no difference was observed between genotypes (Fig. 7A). As Arc and c-Fos upregulation is usually observed in a small subset of hippocampal cells (*43*) while our RNAseq approach examined the entire population of CA1 neurons, we then tested whether quantitative genotype-dependent differences become visible in an IHC-approach visualizing single cells. Consistently, Arc^+^ and c-Fos^+^ cells were increased throughout the hippocampal CA1 in wildtype mice after spatial exploration (p=0.03 for Arc and p=0.01 for c-Fos). However, this immediate early gene response was impaired in mice lacking NEIL3 (p=0.18 for Arc and p=0.06 for c-Fos; p=0.0258 for c-Fos comparing wildtype vs. *Neil3^−/−^* after exploration, two-way ANOVA) (Fig. 7B, C), suggesting a role of NEIL3 in modulating the expression of IEGs in response to spatial experience.

**Figure 7.**
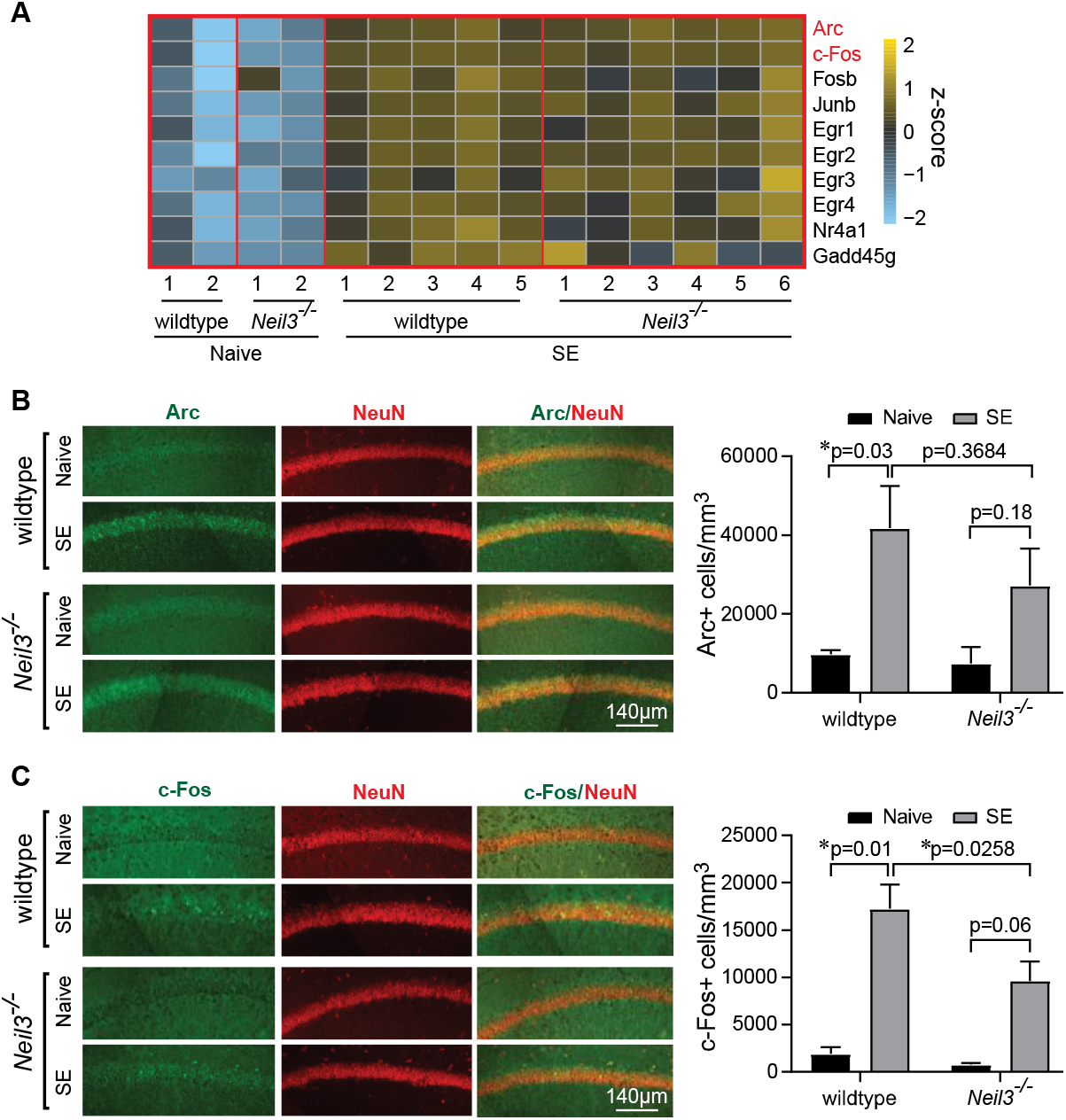
Spatial experience induced expression of immediate early genes in wildtype and *Neil3^−/−^* CA1 neurons. (**A**) Heatmap showing z-score normalized FPKM expression levels of selected immediate early genes (based on whole-CA1 RNAseq) from wt and *Neil3^−/−^* mice without (Naive) or after **<u>s</u>**patial **<u>e</u>**xploration (“SE”). (**B**) Dorsal hippocampal CA1 was immunostained with antibodies against Arc (green) and NeuN (red). The number of Arc positive CA1 neurons in wildtype and *Neil3^−/−^* mice with or without behavior strained was analyzed. (**C**) Similarly, dorsal hippocampal CA1 was immunostained with antibodies against c-Fos (green) and NeuN (red). The number of c-Fos positive CA1 neurons in wildtype and *Neil3^−/−^* mice with or without behavioral intervention was analyzed. Statistics were conducted at animal level (3 animals with 2 hippocampal slices for each genotype, two-way-ANOVA, Sidak’s correction, error bars indicate SEM).

## DISCUSSION

Here, we provide evidence that NEIL3 modulates transcription of genes involved in shaping the functional plasticity of hippocampal place cells. We revealed impaired spatial stability in *Neil3^−/−^* CA1 place cells, demonstrating a functional interference of NEIL3 in CA1 neurons that is associated with impaired spatial cognition. We characterized the contribution of NEIL3 to spatial experience dependent CA1 transcriptome, identifying NEIL3-dependent modulation of gene expression in epigenetic and synaptic regulation. We observed spatial-experience induced changes of NMDAR subunits and immediate early genes in CA1 pyramidal neurons, implicating a role of NEIL3 in the molecular correlates of constrained spatial memory.

### NEIL3 impacts functional plasticity of hippocampal place cells

Hippocampal place cells, as one of the most remarkable neuronal correlates for spatial cognition, have been widely studied to understand the memory mechanisms in the hippocampus. Place cells are selectively activated in a particular location of the environment (*19*) and are able to keep the same firing pattern (“place field”) for days and months, suggesting their encoding for long-term memory of a learned environment (*44*). Meanwhile, place cells often alter their firing patterns in response to environmental changes, suggesting their ability of dissociating the dissimilarities and generating new maps for a novel environment (*24, 25*). In this study, we recorded the activity of neurons in the hippocampal CA1 region of wt and *Neil3^−/−^* mice while the animals were freely moving in an open field environment and examined whether NEIL3 contributes to the spatial plasticity of hippocampal neurons.

Similar numbers of place cells were recorded in wildtype and *Neil3^−/−^* CA1. No significant difference was observed in terms of firing rate, field size and the spatial information content. However, a higher fraction of place cells in *Neil3^−/−^* CA1 displayed multiple environment-specific firing fields (Fig. 1F). *Neil3^−/−^* place cells reliably remapped to the new environment as the wildtype ones (Fig. 2A-B and Fig. S2B), indicating an intact ability of generating new maps. However, an increased fraction of *Neil3^−/−^* place cells kept remapping in the familiar environment when recorded at different trials with a 50min or 24h interval (Fig. 2B/D, Fig. S2D), demonstrating an impaired long-term spatial stability. These findings bear a striking similarity to previous studies using NMDA receptor blockade (*45*). Thus, we postulate that NEIL3 impacts experience-dependent synaptic alterations essential for glutamatergic signaling in CA1 neurons leading to a functional distortion of CA1 place cells to retrieve place maps of a known environment.

Of note, within each recording session, the place fields of *Neil3^−/−^* CA1 neurons were stable and coherent (table in Fig. 1E), implying that *Neil3^−/−^* CA1 place cells reliably maintained the maps once it is selected. Interestingly, preserved within-session spatial stability and frequent remapping across trials were also observed in CA1 place cells of aged animals (*46, 47*), associating with the decreased spatial cognition during aging as observed in the behavior of *Neil3^−/−^* mice (*6*). Future studies should therefore explore NEIL3-dependent mechanisms regulating the age-dependent decline of hippocampal functions.

### NEIL3 modulates the expression of neuronal genes involved in synaptic regulation

Experience-dependent changes in transcription play a pivotal role in the plasticity of neurons and neural circuits for cognitive function and behavior (*14*). The association of NEIL3 with transcriptional modulation has been recently reported. Dysregulated genes related to cardiovascular development and connective tissue disorders were revealed in NEIL3-deficient hearts, whereas no increase in steady state level of oxidative DNA base lesions and no genome-wide hypermutator phenotypes were observed (*48*). Our work identified NEIL3-dependent differentially regulated genes in hippocampal CA1 neurons that are overrepresented in synaptic cellular components and associated with a broad spectrum of neural biological processes, including e.g. synapse organization, synaptic signaling, behavior and neurodevelopment (Fig. 3). Moreover, we pinpointed NEIL3-dependent modulation of gene expression in CA1 neurons specifically after spatial exploration (Fig. 4–5), revealing dysregulated genes involved in synaptic regulation in response to spatial experiences. Importantly, the differential regulation of synaptic factors referred to both excitatory (NMDA-receptor) and inhibitory (GABA-receptor) factors, indicating a function for NEIL3 in maintaining an intact excitatory-inhibitory balance, a disturbance of which has been shown to be crucial for the onset of neurodegenerative diseases (*49*). The Gene Ontology (GO) enrichment analysis provides a thematic classification of high-throughput transcriptome data, positing NEIL3-dependent transcription as central to synaptic homeostasis. Of note, the thematic classification as provided by GO refers to the projected function of the differentially expressed gene, not to the localization where the transcript was isolated. The overall rather subtle transcriptional changes might be due to the behavioral setup we used: a spatial exploration paradigm in the open field leads to neuronal activation, which is less drastic than the learning induced plasticity response.

Differential expression of glutamatergic (NMDA) and GABAergic receptor subunits at *Neil3^−/−^* hippocampus has been observed by IHC in a previous study (*6*). However, at that time, a pan-antibody against both NR2A and NR2B was used and the animals were not exposed to any behavior intervention. The ratio of NR2A and 2B subunits in the forebrain has been linked to memory-related processes, showing an association of an increased NR2A:NR2B ratio with constrained long-term memory (*29–31, 36*). In this study, we observed a tendency of increased NR2B in naive mice lacking NEIL3, whereas the level of NR2A appeared to be reduced. After spatial exploration, a similar upregulation of NR2A, but not NR2B, was detected in both wt and *Neil3^−/−^* CA1. Strikingly, the NR2A:NR2B ratio was significantly increased in *Neil3^−/−^* CA1 after spatial exploration, implicating an impaired long-term spatial memory in NEIL3-deficient mice as described in the previous behavior study (*6*). These findings support the role of NEIL3 in regulation of experience-dependent gene expression in CA1 neurons that are involved in the glutamatergic signaling.

### NEIL3 has a potential impact on neuroepigenetic mechanisms

NEIL3 has been implicated in epigenetic mechanisms involving DNA modifications as well as 3D-genome architecture that are critical for gene regulation (*50, 51*). The precise molecular interplay of NEIL3 with epigenetic marks remains to be elucidated. Here, we find that chromatin-regulating factors such as histone and DNA modifying enzymes are differentially regulated in the absence of NEIL3 (Fig. 5C). For example, downregulation of several histone-lysine-methyltransferases (KMTs), including *Kmt2a* (log2FC=-0.42, Padj=0.002), *Kmt2b* (log2FC=-0.43, Padj=0.001), *Kmt2d* (log2FC=-0.45, Padj=0.003) and *Dot1l/Kmt4* (log2FC=-0.67, Padj=0.006), DNA methyltransferase 3A (*Dnmt3a*, log2FC=-0.41, Padj=0.0003) and TET family methylcytosine dioxygenases, *Tet2* (log2FC=-0.62, Padj=0.005) and *Tet3* (log2FC=-0.40, Padj=0.009), was observed, all of which have been implicated in memory formation (*52–59*). As this study is based on a non-conditional knock-out model, this might reflect an overall adaptation of the organism to the absence of NEIL3. Nonetheless, it points to a potential role of NEIL3 in regulating the chromatin structure and thereby shaping transcription. Our data suggest a role of NEIL3 in mature, postmitotic neurons as a transcription modulator, while previous studies mainly locate NEIL3 function in proliferating, immature cells, referring to its canonical role as a DNA repair enzyme (*5–7*). The NEIL3-dependent epigenetic mechanisms should be further investigated. It is known that learning induces alterations at the level of synaptic physiology as well as chromatin organization (*38, 60, 61*). Our work highlights the impact of NEIL3 in experience-induced epigenomic and synaptic changes, which not only links to the previously described impaired spatial learning and memory phenotype in *Neil3^−/−^* mice (*6*), but also posits NEIL3 as a potential regulator of the epigenomic response to learning and memory formation in general.

### NEIL3 modulates experience-induced immediate early genes in CA1 neurons

Immediate early genes (IEGs) such as *c-fos* and *Arc* are rapidly upregulated in subsets of neurons in response to sensory and behavioral experiences, allowing for the functional adaptation of these neurons and the storage of long-term memory (*37, 38, 61*). Recent studies describe progressively stabilized IEG activation patterns in distinct CA1 neuronal ensembles over repeated visits to the same environment for weeks, supporting that the long-term consolidation of hippocampal plasticity patterns is required for long-term memory formation (*62*). In our spatial experience-dependent transcriptome study, wildtype and *Neil3^−/−^* mice were experienced in the familiar open-field environment 20min/day for 6 days and bulk CA1 samples were collected on the test day 7 after the last trial either the familiar or a novel environment. A 20min exploration allows a thorough and inter-individually equal coverage of the open field environment. This is the same behavior paradigm that we used to record the activity and induce remapping of CA1 place cells (see methods). We observed spatial exploration induced upregulation of several IEGs in wt and *Neil3^−/−^* CA1 (Fig. 7A), which is consistent with the long-term stabilization of IEG activation patterns as previously described (*62*). We did not detect quantitative differences of IEG transcripts between genotypes and no immediate transcriptional changes were observed in CA1 after novel-environment stimuli. As we extracted RNA from a pool of CA1 pyramidal neurons, this could be explained by sparse IEG activation in distinct subpopulation of neurons at CA1, although different environments evoked comparable levels of bulk IEG activity (*43, 62*).

Interestingly, by using immunochemistry and 3D image analysis, we observed that spatial exploration induced activation of Arc and c-Fos was reduced at CA1 lacking NEIL3 (Fig. 7B-C), suggesting that NEIL3 is involved in the maintenance of IEG-dependent ensemble neural plasticity for long-term memory. Arc^+^ and c-Fos^+^ neuronal ensembles in the hippocampus have been explored as engram cells that are associated with memory traces (*63–65*). A recent study revealed that c-Fos^+^ engram cells in CA1 served as an index for contextual memory but not for spatial memory (*66, 67*). The significantly decreased number of c-Fos^+^ cells in *Neil3^−/−^* CA1 after spatial exploration would therefore implicate a potential role of NEIL3 in regulating context-specific memory. An interesting future approach would thus be an activity-dependent memory tagging strategy allowing for the observation, in-and reactivation of memory engrams (*43*) in order to delineate the interplay of NEIL3 and engram formation.

## MATERIALS AND METHODS

### Subjects

3-6 months old wildtype (C57Bl6N) and *Neil3^−/−^* mice (males) with an approximate body weight of 35+/− 3g were used for all experiments. *Neil3^−/−^* mice were generated and described previously (*7*). Animals were housed with their litter-mates in 1717×545×2045mm (LxWxH) cages with free food and water access in a dedicated housing room (temperature 22°C+/− 1°C and humidity 55% +/− 5%) with a 12h light/dark cycle (lights on 7 pm to 7 am). We avoided a priori differences in rack cage position to account for potential differences in light intensity between rack positions. Implanted animals were housed individually after surgery with food and water ad libitum and monitored daily during the experimental period. Behavior and neuronal recording experiments were performed in the dark phase. All experiments were conducted in accordance with the Norwegian Animal Welfare Act and approved by the Norwegian Animal Research Authority (FOTS11659).

### Spatial-exploration and behavioral paradigms

As for assessing the experience-dependent transcriptome, wildtype and *Neil3^−/−^* mice were habituated in the open field of familiar environment (Room A, 50×50cm plastic box with 30cm height of black walls and an A6-sized white cue card fixed at the north side constantly, dim light) with cues and food rewards (crushed Cheerios chocolate loops, Nestle, United Kingdom) for 20 min daily in a sequence of 7 days. The box rested on a table at a height of 100cm and surrounded by black curtains and the animal entered the environment from a constant side. After every exposition to the spatial exploration paradigm, the open field area was cleaned using perfume-free soapy water (antibac skincare, Asker, Norway). On the test day (Day 7), half of the animals from each genotype were exposed in the familiar environment for 20 min, then rested for another 20 min before termination. The other half of the animals from each genotype were tested in a novel open field environment (Room B, 50×50cm plastic box with 30cm height of white walls and an A6-sized black cue card fixed at the north side constantly, dim light) for 20 min, then rested for another 20 min before termination. As for the IHC studies, wildtype and *Neil3^−/−^* mice were only exposed to the familiar open field environment (20 min/training, 7 days) and terminated 40 min after the last test in Day7. As for the extracellular neuronal recording experiments, animals were habituated and recorded in the familiar open field environment in Room A and tested in a novel open field environment in Room B (same setup as previous).

### Laser dissection of CA1-specific brain tissue

The brain was taken out from the mouse without intracardial perfusion, mounted on a cryostat metal socket (Leica CM3050S, Leica, Nussloch, Germany) using a drop of mounting media (Tissue-Tek OCT compound, Sakura Fintek, Torrance, CA, USA), immediately frozen using pulverized dry ice (101 Cold Spray; Taerosol, Kangasala, Finland) and kept in liquid nitrogen before frozen sectioning. A thickness of 8μm coronal brain sections were collected and mounted immediately on membrane slides (Molecular Machines and Industries GmbH, Eching, Germany). Typically, we mounted 5-6 slices per slide and collected a total of 20 slides for tissue isolation per animal, which usually comprised the entire rostro-caudal axis of the hippocampal formation. Brain slices were stained using a modified cresyl violet fast-staining procedure, including sequential steps (***i***) dehydration (7 dips each in 70%, 80%, 90%, 100% EtOH), (***ii***) tissue clearance in Xylene (Sigma-Aldrich, Oslo, Norway) for 1min, (***iii***) rehydration in cresyl violet solution (Sigma-Aldrich, Oslo, Norway) for 90s without motion, (***iv***) final dehydration and clearance in xylene, (***v***) drying for 15-20min. Afterwards, CA1 dissectates were collected using a laser dissection microscope (Molecular Machines and Industries, CellCut on Olympus IX71, Eching, Germany). The hippocampal CA1 area was identified using the Allen Mouse Brain Atlas (Allen Brain Institute) as a reference. For each slice, the whole CA1 area was defined manually (see Figure 1A). We collected 20 CA1 dissectates in one isolation cap (Molecular Machines and Industries GmbH, Eching, Germany) and finished one animal within the day. CA1 samples were lysed in RLT lysis buffer (AllPrep Kit, Qiagen, Hilden, Germany) and frozen at −80°C until further processing.

### RNAseq and quality validation

Laser-dissected CA1 samples were lysed in RLT lysis buffer (AllPrep Kit, Qiagen, Hilden, Germany) using a bead homogenizer (MagNA lyser, Roche Diagnostics, Oslo, Norway). RNA was extracted using either the AllPrep Kit or the RNeasy Mini Kit from Qiagen according to the manufacturer’s instructions. RNA samples typically yielded >100ng of RNA with a RIN value of > 7 as determined by Bioanalyzer (Agilent Technologies, Santa Clara, USA).

Whole-transcriptome sequencing was done by BGI Group (BGI Genomics Co., Ltd., Hong Kong, China) according to their in-house protocol. In brief, after quality control using the Agilent 2100 Bio analyzer (Agilent RNA 6000 Nano Kit, Agilent Technologies, Santa Clara, CA, USA), poly-T oligo-attached magnetic beads were used to purify poly-A containing mRNA. After mRNA-fragmentation (divalent cations, high temperature), RNA was copied into cDNA using reverse transcriptase and random primers with subsequent second strand synthesis using DNA Polymerase I rand RNase H. Qubit quantification (Thermo Fisher Scientific, Waltham, MA, USA) was used to quantify the PCR yield after amplification. The final library was constructed making a single strand DNA circle. Rolling circle replication created DNA nanoballs, which yields more fluorescent signal during sequencing. Pair-end reads of 100bp were read via the BGISEQ-500 platform.

The subsequent bioinformatic analysis was in part done by BGI (BGI Genomics Co., Ltd., Hong Kong, China) using the following workflow: In a first step, low quality reads were filtered using the BGI-internal software SOAPnuke (reads with adaptors, reads in which unknown bases are >10% and low-quality reads with threshold 15 were removed. Genome mapping was done using the HISAT software. Transcripts were then reconstructed using StringTie and compared to reference using Cuffcompare. For gene expression analysis, clean reads were mapped to reference using Bowtie2 and then, gene expression levels were calculated using the RSEM software package.

### Analysis of differential gene expression

Based on the sequencing results from BGI (un-normalized estimated counts) the count matrix values were generated, and the metadata matrix was prepared. For the exploratory analysis, the counts were transformed with regularized logarithm using rlog function and the PCA was then performed. The differential gene expression analysis based on binomial distribution was performed in R ver. 3.6.2, Platform: i386-w64-mingw32/i386 (R Core Team, 2019), Bioconductor ver. 3.10 (*68*) using DESeq2 ver. 1.26.0 (*27*) and plots were generated in ggplot2 (*69*). Factor levels were set using contrast function as Mutant vs. Wild Type. The alpha parameter was set to 0.05 and adjusted p value was calculated using Benjamini-Hochberg correction for multiple testing. To further analyze the differentially expressed genes (DEGs) the list was made with ABS (log2 Fold Change) threshold set to 0.3. To visualize the results volcano plot was generated using EnhancedVolcano (*70*) and heatmaps of FPMK values were generated with pHeatmap (Kolde, 2019).

For the over-representation analysis the online version of PANTHER Classification System release 15.0 (Released 2020-07-28) was used (*28*). To perform the over-representation test (using Binomial test and Bonferroni correction), the list of DEGs was uploaded and the mouse genome was set as background. The test was performed for each functional category (GO Ontology database released 2020-09-10): Biological Process, Cellular Compartment and PANTHER Protein Class.

### Surgical Procedure

Mice were anesthetized using isoflurane (induction chamber level of 3.0% with an air flow at 3000 ml/min, gradually reduced after the mice were secured in the stereotaxic apparatus to 0.5-1% isoflurane with an air flow at 2000 ml/min). Anesthesia was upheld via constant isoflurane application throughout the surgical procedure, using the physiological breathing rate of mice as a guidance for controlling anesthesia depth. A weight-adapted dose of buprenorphine (Temgesic, Indivior, Dublin, Ireland) was given either intraperitoneally or subcutaneously at least 15min prior to the first incision. A local dose of bupivacaine (Marcain, Aspen, Ballerup, Denmark) was injected subcutaneously in the incision area of the mouse scalp ca. 5min prior to the first incision. A prophylactic, weight-adapted dose of meloxicam (Metacam, Boehringer-Ingelheim Vetmedica, Copenhagen, Denmark) was given 15min before the end of the surgical procedure. The total anesthesia time was typically around 60min (45min-90min). Throughout surgery, eyes were protected using moisturizing eye cream (ViscoTears, Thea Pharma, Wetteren, Belgium). Post-operative pain was controlled using buprenorphine and/or meloxicam according to the animal’s need as assessed by a combined score of facial grimaces, behavior, fur state and weight. Mice generally showed mild to moderate abnormalities in these categories for a duration of 2-3 days and reached completely normal levels no later than one week after the procedure.

A microdrive with an assembly of four tetrodes (sixteen electrodes) was inserted into the right hemisphere above the CA1 area of the dorsal hippocampus (stereotactic coordinates: AP[Bregma]: 2mm, ML: 1.8mm, DV: 0.8mm). The electrodes were made of 17-mm polyimidecoated platinum-iridium (90 to 10%) wire (California Fine Wire Co., USA) and plated with platinum (Platinum Black Plating Solution, Neuralynx, Inc., USA) to reduce electrode impedances to ~200 kWat 1 kHz. A high-speed drill (Model1474, David Kopf Instruments, Tujunga, CA, USA) was used to penetrate the skull right above the area of tetrode insertion. Using a 27G needle (Sterican, B.Braun, Melsungen, Germany), the dura was gently nicked but not removed to allow the tetrode entering the neurocranium with ease. A metal 19G metal cannula was placed around the tetrode, resting loosely on the dura. The area of bone removal was kept moist using dental sponge (Spongostan Dental, Ethicon, Bridgewater, NJ, USA) soaked in saline 0.9% (B.Braun, Melsungen, Germany). A small rest of dental sponge was left in place after the surgical procedure, helping to minimize the freedom of movement of the cannula-tetrode entity. A jeweler’s screw was fixed to the skull serving as a ground electrode. The Microdrive and ground screw were fixed to the skull using dental cement. After recovery from the surgical procedure, neuronal activity was recorded while the mouse was freely moving in an open field environment.

### Neuronal Recording Procedures

Animals were exposed to an open field environment (50×50cm plastic box with 30cm height of black walls and an A6-sized white cue card fixed at the north side constantly, dim light) 3 days after implantation and neuronal activities were recorded one week later when tetrode turning was started. The implanted microdrive was connected to the recording equipment (Axona, St. Albans, UK) via AC-coupled unity-gain operational amplifiers, using a counterbalanced cable that allowed the animal to move freely in the recording box. Recording data from HPC were collected using DacqUSB software from Axona. Recorded signals were amplified 8000 to 25,000 times and band-pass filtered between 0.8 and 6.7 kHz. The recording system tracked the position of two light-emitting diodes (infrared LEDs, one large and one small, 5 cm length) on the head stage (weight 5.12g) using an overhead video camera.

Neuronal activity was recorded while the animal was free to move in the recording box for a duration of 20 min. The behavior of mice was motivated by crumbs of chocolate loops (Cheerios, Nestle, United Kingdom) randomly scattering in the open field area. The tetrodes were lowered in steps of 25 mm until single neurons could be isolated at appropriate depths. When putative place cells were observed, the global remapping of place cells was monitored following five sequential recording sessions in the familiar and novel open field environment (Fig. 2A). The familiar environment (50×50cm square box, black walls with a white cue card) is in room A where animals were habituated. The novel environment is white square boxes (50×50cm) with a black cue card in a different room compartment (named room B). In between recording sessions, mice were rest in a 30×20cm transparent plastic box (similar as their home cage) for about 3 min. After every session, the open field area was cleaned using perfume-free soapy water (antibac skincare, Asker, Norway). After collection of each data set, the tetrodes were moved further until new well-separated place cells were encountered. In general, CA1 place cells were recorded in a DV-depth of 0.8-1.4 mm.

### Spike Sorting and Rate Maps

The offline graphical cluster-cutting software Tint (Axona Ltd., St. Albans, United Kingdom) was used for spike sorting. The software provided a multidimensional parameter space consisting of two-dimensional projections with spike wave amplitudes and waveform energies, which were used to sort spikes and identify spike clusters belonging to one cell. Autocorrelation and cross-correlation functions were used as additional criteria to separate individual cell clusters reliably. Animal’s position data were estimated based on the tracking of the two LEDs connecting to the head stage and the microdrive. All data were speed filtered, including spikes only when the animal had an instantaneous running speed of 2-100 cm/s. As described previously (*71*), the distribution of firing rates was determined by the number of spikes as well as the time spent in each bin (2.5×2.5 cm) of the recording area (50×50cm). A 21-sample boxcar window filter (400ms, 10 samples on each side) was used for smoothing the recorded path. Additional smoothing using a quasi-Gaussian kernel over the surrounding 5 × 5 bins was applied to generate maps of spike number and time. The quotient of spike number and time for each bin of the smoothed map was used for defining the firing rate, with the bin showing the highest rate defined as the “peak rate”.

### Analysis of place cells

Place cells were defined by comparing each cell’s spatial information score with the distribution of information scores for rate maps generated from randomly shuffled data as previously described (*23*). The spatial information content in bits per spike was calculated as

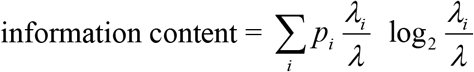

where *λ*_*i*_ is the mean firing rate of a unit in the *i-*th bin, *λ* is the overall mean firing rate, and *p*i is the probability of the animal being in the *i-*th bin (occupancy in the *i-*th bin / total recording time) (*72*). The raw data were first divided into quadratic bins (2.5×2.5 cm). Then the firing rate at each point in the environment was estimated by expanding a circle around the point until

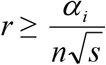

where *r* is the radius of the circle in bins, *n* is the number of occupancy samples within the circle, *s* is the total number of spikes in those occupancy samples, and the constant *α* is set to 10000. With a position sampling rate of 50 Hz, the firing rate at that point was then set to 50 · *s*/*n*.

The chance level for spatial information was determined by a random shuffling procedure. The shuffled data are generated by time-shifting the entire sequence of spikes fired by one cell along the path of the animal: Each time shift equaled a random interval of a minimum of 20s and a maximum of the entire trial length minus 20s (in our case 1200s – 20s = 1180s), with the end of each trial wrapped to the beginning. The shuffling procedure was repeated 100 times for each of the 419 cells recorded in the wildtype mice, yielding a total of 41900 permutations and each of 402 cells recorded in the *Neil3^−/−^* mice, yielding a total of 40200 permutations. A rate map was generated for each permutation and the distribution of spatial information values across all permutations of all cells was determined. The 95th percentile (see Figure 5) was used as a threshold to define place cells.

A place field was defined as an area equaling/larger than 50 cm^2^ (8 or more 2.5 cm × 2.5 cm bins) where the firing rate was above 20% of the peak rate. Only place fields with a peak firing rate of at least 1Hz as well as a minimum of 100 spikes were included in the analysis. Interneurons and bypassing axons were defined by an average peak-to-trough waveform duration of less than 200μs and excluded from the analysis. Spatial coherence was estimated as the first-order spatial autocorrelation of the smoothed rate maps, i.e. the mean correlation between firing rate of each bin and the averaged firing rate in the eight adjacent bins (*73*). In-session spatial stability was calculated by the spatial correlation of place field maps from the first and second half of the open field trial.

The spatial correlation between trials was determined for individual cells as previously described (*23, 26*). In brief, the rates of firing in corresponding bins of smoothed rate maps were correlated with one another, leading to a correlation procedure containing both the localization and number of spikes as the core variables. Correlation coefficients were calculated based on the full trial. Only place cells passing the 95th percentile criteria were included in the analysis.

### Histology and reconstruction of recording positions

Electrodes were not moved after the final recording session. Mice were terminated using a combination of isoflurane (Baxter, Oslo, Noway) and pentobarbital (>200mg/kg bodyweight) and then transcardially perfused with saline 0.9% followed by 4% paraformaldehyde. The electrodes were turned all the way up before the brain was extracted. Brains were quickly frozen and sectioned by a cryostat (Leica CM3050S, Nussloch, Germany, tempered to −25°C) at a thickness of 30μm in the sagittal plane. All sections around the area of the tetrode trace were collected and mounted on histological glass slides (SuperFrost, Menzel, Braunschweig, Germany). Slides were left to dry at room temperature overnight, stained using a standard cresyl violet staining protocol (see paragraph on laser dissection) and imaged.

### Immunohistochemistry

Sagittal brain sections (30 μm) were collected by frozen sectioning. Sections were incubated in the antigen retrieval buffer (40mM trisodium citrate, pH6.0) at 99°C for 3 min. After washing in PBS, sections were preincubated in the blocking buffer (PBS with 5% normal goat serum, 1% BSA, and 0.1% Triton X-100) at 4° for 2 hours and then incubated with the diluted primary antibodies in dilution buffer (PBS with 1% normal goat serum, 1% BSA, and 0.1% Triton X-100) at 4° overnight. The next day, sections were washed in 1xPBST 3 times at room temperature and incubated with the diluted secondary antibodies at room temperature for 2 hours. After 3 times washing in 1x PBST, sections were mounted onto glass slides and dry overnight. Lastly, sections were incubated with Dapi (1μg/mL in PBS), washed and cover-slipped with mounting oil (ProLong™ Gold Antifade Mountant with DAPI, ThermoFisher). As details for primary and secondary antibodies, see the key material table.

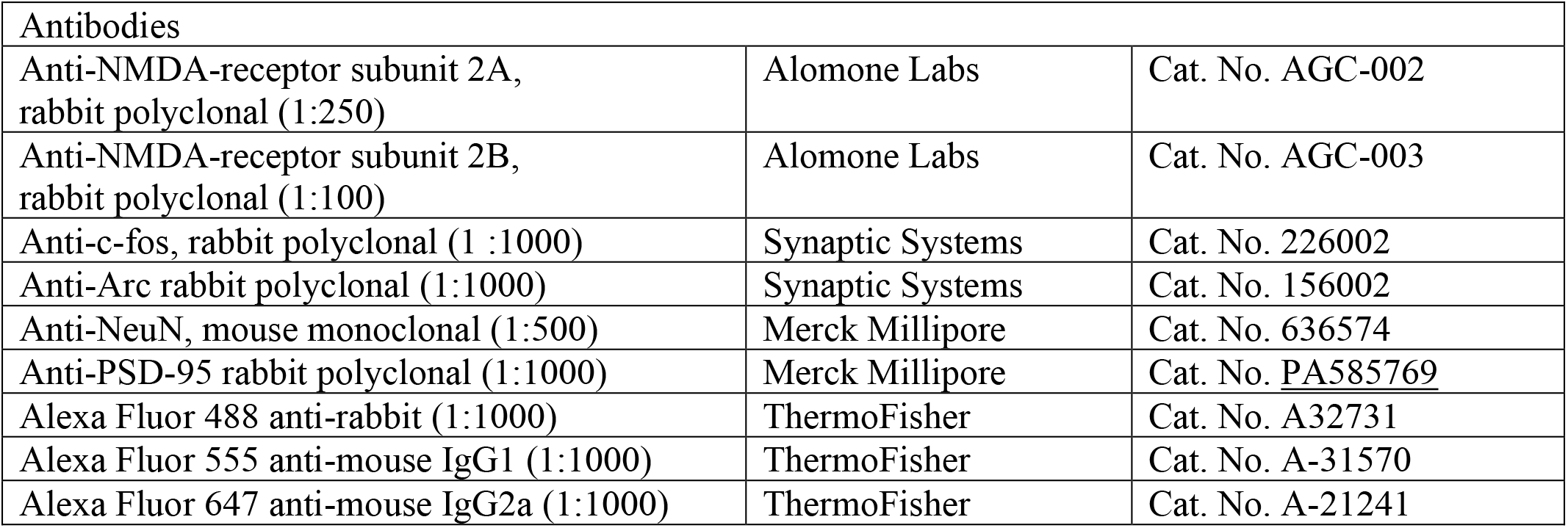

| Antibodies |  |  |
| --- | --- | --- |
| Anti-NMDA-receptor subunit 2A, rabbit polyclonal (1:250) | Alomone Labs | Cat. No. AGC-002 |
| Anti-NMDA-receptor subunit 2B, rabbit polyclonal (1:100) | Alomone Labs | Cat. No. AGC-003 |
| Anti-c-fos, rabbit polyclonal (1 :1000) | Synaptic Systems | Cat. No. 226002 |
| Anti-Arc rabbit polyclonal (1:1000) | Synaptic Systems | Cat. No. 156002 |
| Anti-NeuN, mouse monoclonal (1:500) | Merck Millipore | Cat. No. 636574 |
| Anti-PSD-95 rabbit polyclonal (1:1000) | Merck Millipore | Cat. No. <u>PA585769</u> |
| Alexa Fluor 488 anti-rabbit (1:1000) | ThermoFisher | Cat. No. A32731 |
| Alexa Fluor 555 anti-mouse IgG1 (1:1000) | ThermoFisher | Cat. No. A-31570 |
| Alexa Fluor 647 anti-mouse IgG2a (1:1000) | ThermoFisher | Cat. No. A-21241 |

### Confocal imaging and 3D image analysis

All fluorescent images were taken by confocal microscope (Zeiss LSM880). For synaptic markers, a Plan-Apochromat 40x/1.4 Oil DIC M27 objective (Carl Zeiss, Jena, Germany) was used. The size of the image window was 700×700μm (x/y 2000pixels of 0.35μm) and a z-interval of 0.5μm was applied. The distal and proximal part of the CA1 was imaged based on NeuN and DAPI stainings as an anatomical orientation (center of proximal square set at 0.25x total CA1 length measured from proximal end, center of distal square at 0.75xtotal CA1 length). For immediate early gene immunohistochemistry, a Plan-Apochromat 20x/0.8 M27 (Carl Zeiss, Jena, Germany) objective was used, a z-plane interval of 2μm was deemed sufficient to identify whole cells. The entire CA1 region was imaged as a region of interest based on NeuN and DAPI staining as an anatomical orientation.

Imaris 9.3 (Bitplane, Zurich, Switzerland) was used to analyze immunoreactivity for synaptic markers and immediate early genes. As for the study of synaptic markers, a 3D reconstruction of the whole z-plane dataset was created. The strata pyramidale/oriens/radiatum were identified as regions of interest using the “surface” tool in Imaris and copied to every z-plane accordingly. Based on the surface selection, a 3D-frame was created and the parameter of interest “masked” according to this frame. The “Spots Wizard” in Imaris was used to model areas of synaptic reactivity within this masked channel (1μm spot diameter for bouts of synaptic reactivity, 5μm diameter for IEGs; background subtraction used in all cases). After visual inspection of the modelled signal for biological relevance, the threshold for Quality Filter (see bitplane.com/imaris) was defined and kept throughout the analysis. Background subtraction was done for every specimen analyzed to account for intensity variations despite identical immunohistochemistry and confocal parameters. 2/3 of the region of interest had to be intact (i.e. not damaged by tissue cracks, covered by imaging artefacts etc.) to be included in the analysis. As for the study of immediate early genes, c-Fos and Arc positive cells in CA1 were identified using the tool of “Spots Wizard” in Imaris. A spot size of 5μm diameter was selected to spot individual cells. Since the whole-region scans such as the CA1-area for IEGs typically involve variations in signal intensity, the quality measure was manually adjusted when necessary.

### Quantification and statistical analysis

Whenever suitable, a two-way-ANOVA was used to compare results between groups, considering the factors *genotype (wildtype vs. *Neil3^−/−^*) and *intervention (baseline vs. spatial exploration). A correction for multiple testing was performed in all cases, generally employing Sidak’s method. Whenever only one variable differed between groups, a Student’s t-test with Welch’s correction was used. With respect to the previously described nested data problem (*74*), all comparisons were conducted with 1 animal equaling 1 statistical unit. In general, this was done by averaging values of several samples (e.g. several hippocampal slices) taken from the same specimen. The number of statistical units per experiment were as follows: (***i***) extracellular live recordings (Fig. 1–2), n=4 per genotype; (***ii***) transcriptome baseline (Fig. 3), n=2 per genotype; (***iii***) transcriptome “spatial exploration” condition (Fig. 4–5), n=5 for wildtype, n=6 for *Neil3^−/−^*; (***iv***) immunohistochemistry (Fig. 6–7), n=3 per genotype both at “Naive” and “SE”. For the analysis of place cells, a population-based (1 cell = 1 statistical unit) analysis was applied, but only reported results as significant that passed a p<0.05 in a nested t-test (1 animal = 1 statistical unit). All statistical analysis was conducted using GraphPad Prism Version 8.

## Data and materials availability

This study did not generate new unique reagents. All RNA sequencing data of this study have been deposited for public access in the NIH database Gene Expression Omnibus (GEO). Accession codes: GSE148408

## ACKNOWLEDGEMENTS

We are grateful to Edvard Moser and May-Britt Moser for sharing the MATLAB code of place cell analysis and the access of NORBRAIN infrastructures in Trondheim, inspiring discussion, and critical reading of the manuscript. We appreciate feedback on the manuscript provided by Carol A. Barnes, Katja Scheffler and Barbara van Loon. We thank Menno Witter for inspiring discussion and technical support on neuroanatomy, Pål Sætrom for his critical thoughts in RNAseq analysis, Atle van Beelen Granlund and Bjørn Munkvold for sharing the laser dissection and cryostat equipment. This work was funded by Research Council of Norway (FRIPRO 297037 and 287911) and Health Authority Central Norway (Helse Midt-Norge, HMN 46060921).

## AUTHOR CONTRIBUTIONS

N.K. and J.Y. designed the experiments and contributed to data analysis. N.K. conducted all experiments. M.S.F.B. assisted on IHC studies. A.M.B. analyzed the RNAseq data. P.B. contributed to extracellular recording of CA1 neurons. J.Y. and M.B. designed and supervised the research. N.K. and J.Y. wrote the manuscript. M.B. and J.Y. acquired the funding. The authors declare no conflicting interests.

## SUPPLEMENTAL INFORMATION

See separate pdf-file “Supplemental Information.pdf”

## Notes

### Competing Interest Statement

The authors have declared no competing interest.

