## Supplement information for "DNA repair enzyme NEIL3 enables a stable neural representation of space by shaping transcription in hippocampal neurons"

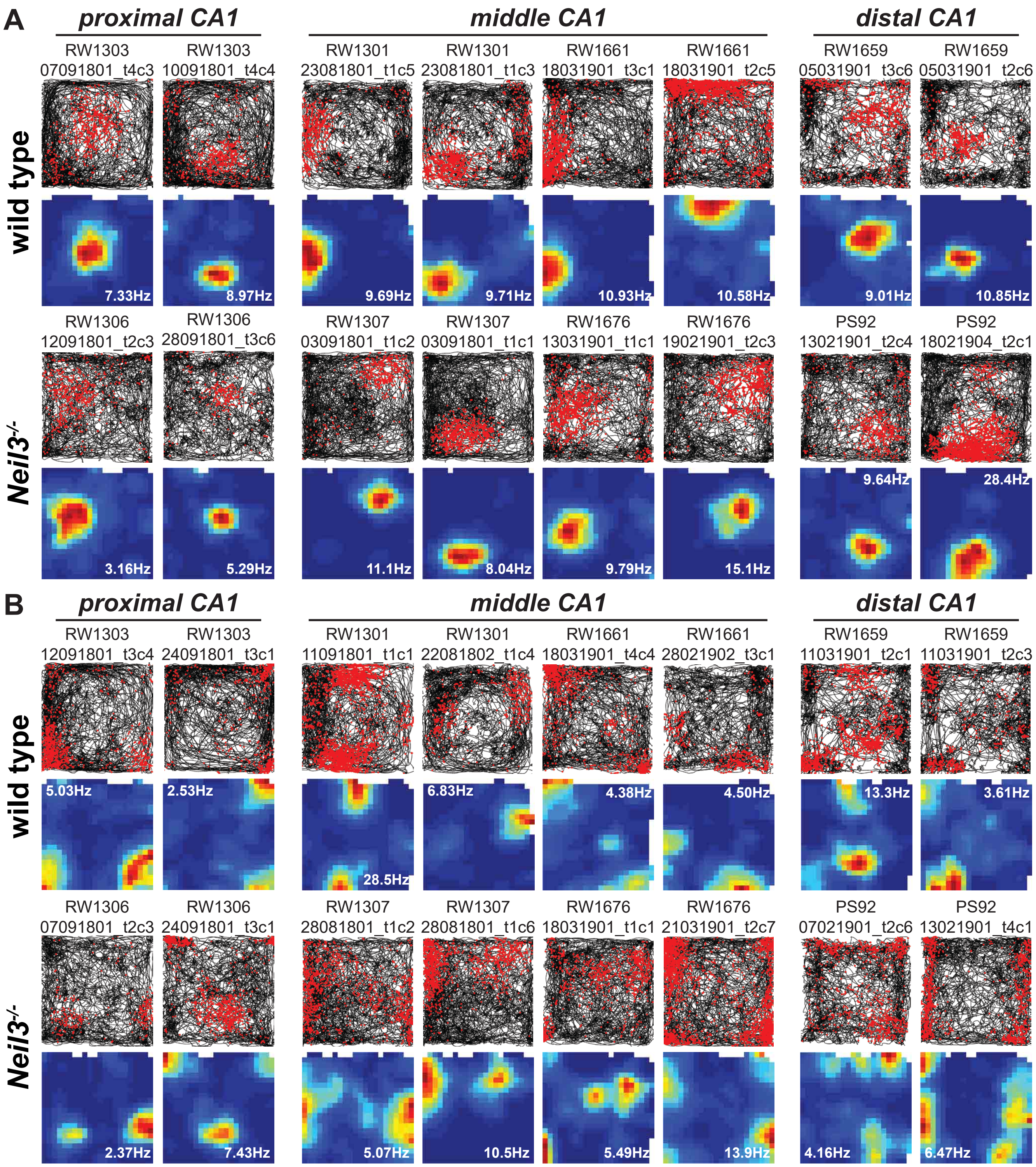


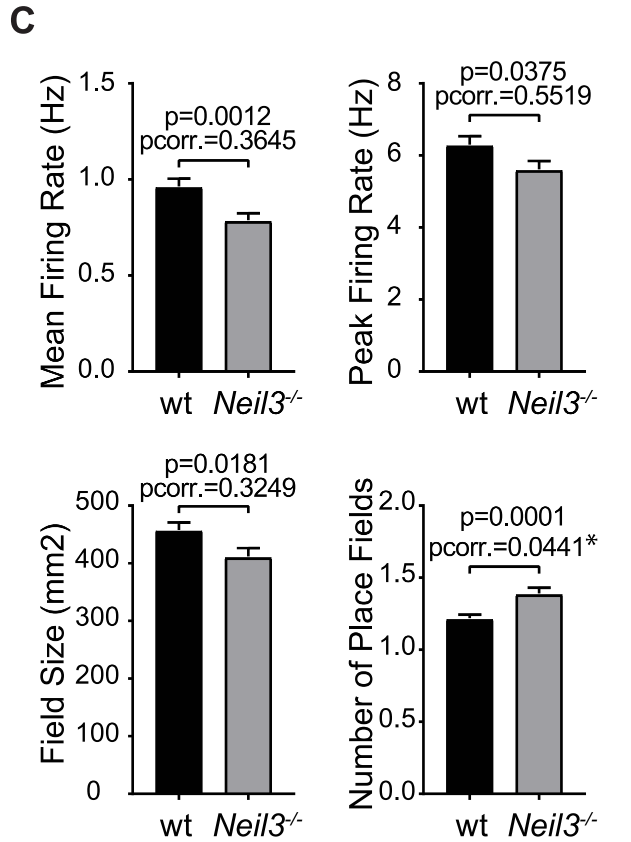


**Supplement Figure 1. Place cells recorded in the CA1 of wildtype and *Neil3*$^{-/-}$ mice.** (**A**) Representative examples of trajectory maps with superimposed spikes (rows 1 and 3, trajectory in black, spikes in red) and color-coded rate maps of single-field place cell activity (rows 2 and 4, red color representing peak firing rate, blue color no firing). Separate images for proximal, middle and distal CA1 are shown. (**B**) Representative examples of place cells with multiple place fields shown as trajectory maps and rate maps as in A. (**C**) Bar plots showing the difference in terms of mean firing rate, peak firing rate, the number and size of place fields of all CA1 place cells recorded in wildtype and *Neil3*$^{-/-}$ mice. Statistics were conducted using unpaired t-test with Welch's correction, at the population level (each cell as statistical unit) and at an animal level using nested t-test (pcorr., each animal equals one statistical unit).

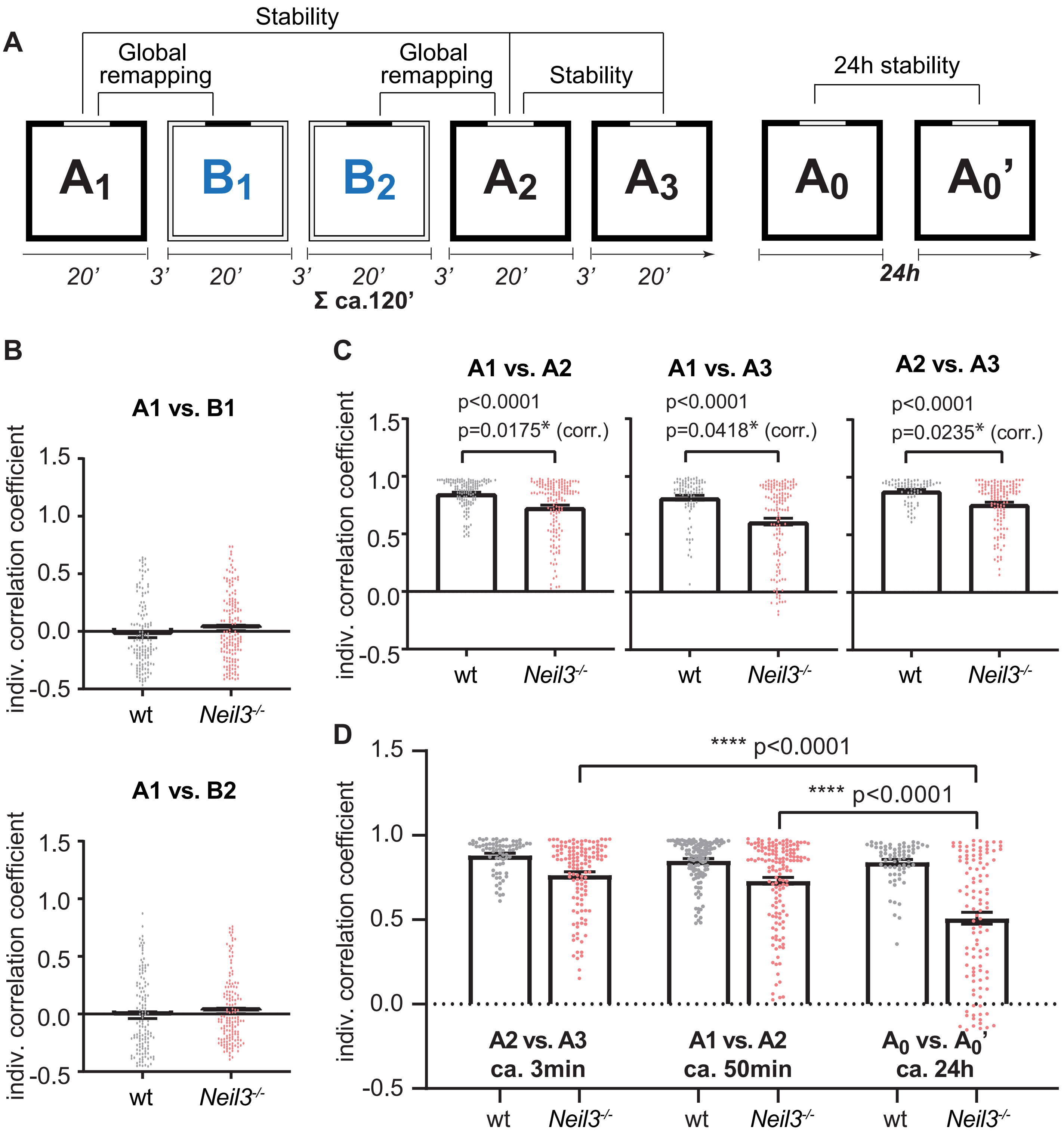


**Supplement Figure 2. Functional plasticity of CA1 place cells in wildtype and Neil3-/- mice.** (**A**) Overview of the experimental set-up. Room A (black wall, white cue card) represents the known environment, Room B (white wall, black cue card) the novel environment. Exploration time for each room was 20 minutes, interceded by a 3 minutes break in a neutral plastic pot. Mice were re-tested in the known environment A after 24h (day 1 A0, day 2 A0'). (**B**) Comparison of spatial cross-correlation coefficients between the known environment A and the first (B1) and second time (B2) the mice were exposed to the novel environment. (**C**) Correlation coefficients for the first (A1 vs. A2, p<0.0001 at a population level/p=0.0175 nested t-test for the comparison wildtype vs. *Neil3*$^{-/-}$ mice) and second (A1 vs. A3, p<0.0001 population/p=0.0418 nested t-test) re-exposure to the known environment in wildtype and *Neil3*$^{-/-}$ mice. (**D**) Gradual deterioration of remapping accuracy in *Neil3*$^{-/-}$ mice over time as compared to wildtype (p<0.0001 for both within-genotype comparisons, mixed-model/Sidak's multiple comparisons test, also see figure 2E).

Figure S3-1

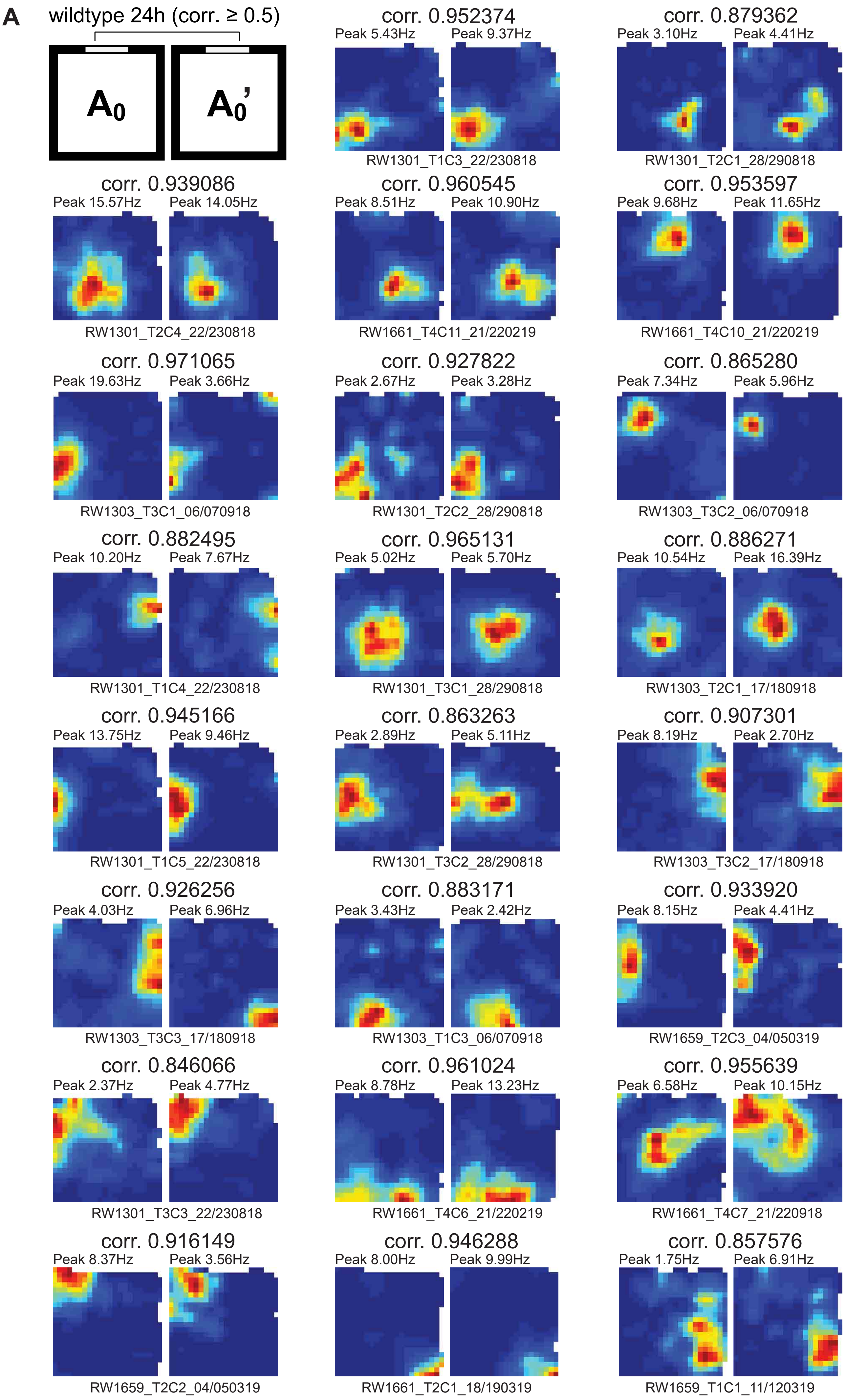
A
wildtype 24h (corr. ≥ 0.5)
A0
A0'
corr. 0.952374
Peak 5.43Hz
Peak 9.37Hz
RW1301_T1C3_22/230818
corr. 0.879362
Peak 3.10Hz
Peak 4.41Hz
RW1301_T2C1_28/290818
corr. 0.939086
Peak 15.57Hz
Peak 14.05Hz
RW1301_T2C4_22/230818
corr. 0.960545
Peak 8.51Hz
Peak 10.90Hz
RW1661_T4C11_21/220219
corr. 0.953597
Peak 9.68Hz
Peak 11.65Hz
RW1661_T4C10_21/220219
corr. 0.971065
Peak 19.63Hz
Peak 3.66Hz
RW1303_T3C1_06/070918
corr. 0.927822
Peak 2.67Hz
Peak 3.28Hz
RW1301_T2C2_28/290818
corr. 0.865280
Peak 7.34Hz
Peak 5.96Hz
RW1303_T3C2_06/070918
corr. 0.882495
Peak 10.20Hz
Peak 7.67Hz
RW1301_T1C4_22/230818
corr. 0.965131
Peak 5.02Hz
Peak 5.70Hz
RW1301_T3C1_28/290818
corr. 0.886271
Peak 10.54Hz
Peak 16.39Hz
RW1303_T2C1_17/180918
corr. 0.945166
Peak 13.75Hz
Peak 9.46Hz
RW1301_T1C5_22/230818
corr. 0.863263
Peak 2.89Hz
Peak 5.11Hz
RW1301_T3C2_28/290818
corr. 0.907301
Peak 8.19Hz
Peak 2.70Hz
RW1303_T3C2_17/180918
corr. 0.926256
Peak 4.03Hz
Peak 6.96Hz
RW1303_T3C3_17/180918
corr. 0.883171
Peak 3.43Hz
Peak 2.42Hz
RW1303_T1C3_06/070918
corr. 0.933920
Peak 8.15Hz
Peak 4.41Hz
RW1659_T2C3_04/050319
corr. 0.846066
Peak 2.37Hz
Peak 4.77Hz
RW1301_T3C3_22/230818
corr. 0.961024
Peak 8.78Hz
Peak 13.23Hz
RW1661_T4C6_21/220219
corr. 0.955639
Peak 6.58Hz
Peak 10.15Hz
RW1661_T4C7_21/220918
corr. 0.916149
Peak 8.37Hz
Peak 3.56Hz
RW1659_T2C2_04/050319
corr. 0.946288
Peak 8.00Hz
Peak 9.99Hz
RW1661_T2C1_18/190319
corr. 0.857576
Peak 1.75Hz
Peak 6.91Hz
RW1659_T1C1_11/120319

**Figure S3-2**

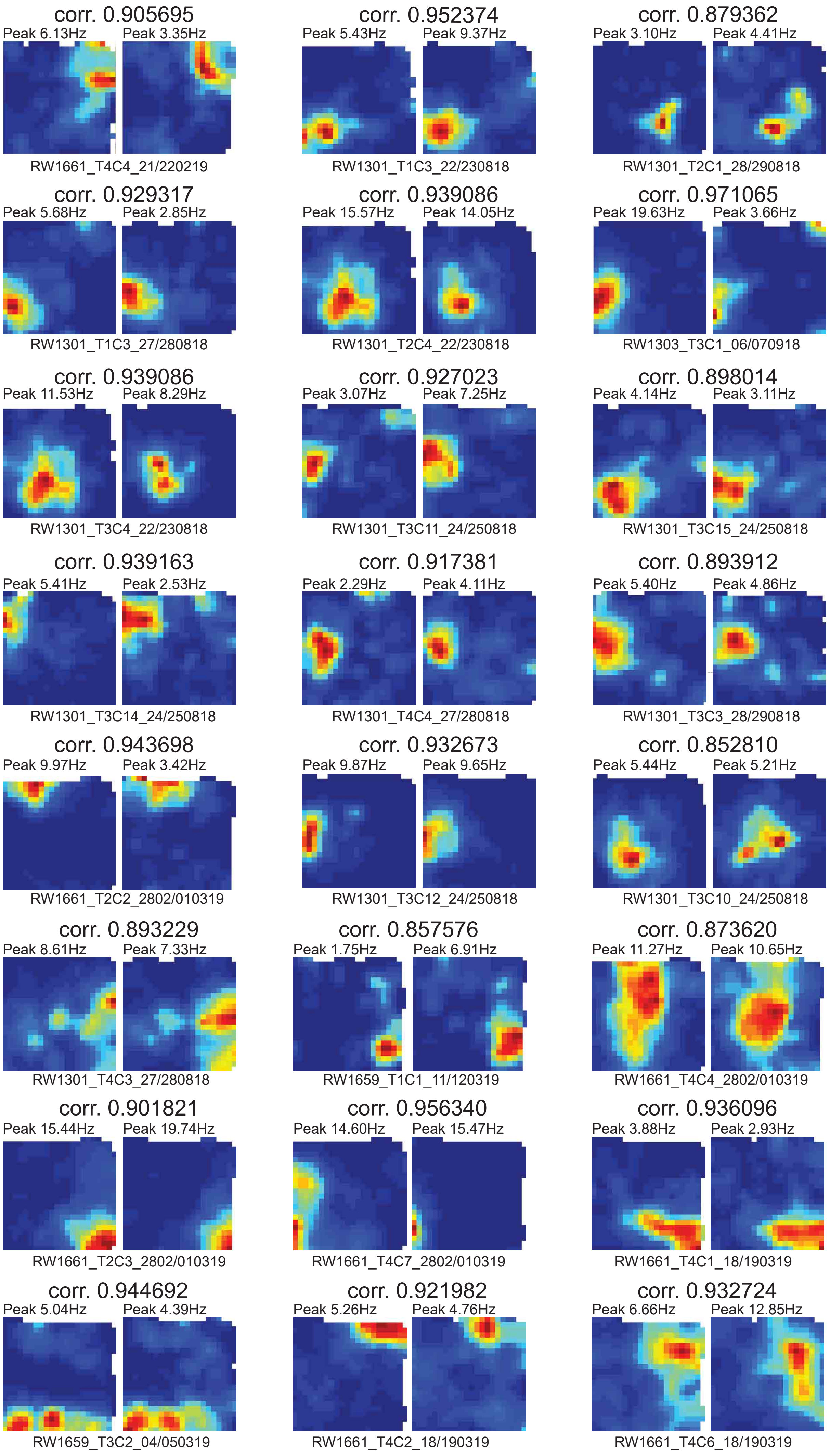
corr. 0.905695
Peak 6.13Hz
Peak 3.35Hz
RW1661_T4C4_21/220219
corr. 0.952374
Peak 5.43Hz
Peak 9.37Hz
RW1301_T1C3_22/230818
corr. 0.879362
Peak 3.10Hz
Peak 4.41Hz
RW1301_T2C1_28/290818
corr. 0.929317
Peak 5.68Hz
Peak 2.85Hz
RW1301_T1C3_27/280818
corr. 0.939086
Peak 15.57Hz
Peak 14.05Hz
RW1301_T2C4_22/230818
corr. 0.971065
Peak 19.63Hz
Peak 3.66Hz
RW1303_T3C1_06/070918
corr. 0.939086
Peak 11.53Hz
Peak 8.29Hz
RW1301_T3C4_22/230818
corr. 0.927023
Peak 3.07Hz
Peak 7.25Hz
RW1301_T3C11_24/250818
corr. 0.898014
Peak 4.14Hz
Peak 3.11Hz
RW1301_T3C15_24/250818
corr. 0.939163
Peak 5.41Hz
Peak 2.53Hz
RW1301_T3C14_24/250818
corr. 0.917381
Peak 2.29Hz
Peak 4.11Hz
RW1301_T4C4_27/280818
corr. 0.893912
Peak 5.40Hz
Peak 4.86Hz
RW1301_T3C3_28/290818
corr. 0.943698
Peak 9.97Hz
Peak 3.42Hz
RW1661_T2C2_2802/010319
corr. 0.932673
Peak 9.87Hz
Peak 9.65Hz
RW1301_T3C12_24/250818
corr. 0.852810
Peak 5.44Hz
Peak 5.21Hz
RW1301_T3C10_24/250818
corr. 0.893229
Peak 8.61Hz
Peak 7.33Hz
RW1301_T4C3_27/280818
corr. 0.857576
Peak 1.75Hz
Peak 6.91Hz
RW1659_T1C1_11/120319
corr. 0.873620
Peak 11.27Hz
Peak 10.65Hz
RW1661_T4C4_2802/010319
corr. 0.901821
Peak 15.44Hz
Peak 19.74Hz
RW1661_T2C3_2802/010319
corr. 0.956340
Peak 14.60Hz
Peak 15.47Hz
RW1661_T4C7_2802/010319
corr. 0.936096
Peak 3.88Hz
Peak 2.93Hz
RW1661_T4C1_18/190319
corr. 0.944692
Peak 5.04Hz
Peak 4.39Hz
RW1659_T3C2_04/050319
corr. 0.921982
Peak 5.26Hz
Peak 4.76Hz
RW1661_T4C2_18/190319
corr. 0.932724
Peak 6.66Hz
Peak 12.85Hz
RW1661_T4C6_18/190319

**Figure S3-3**

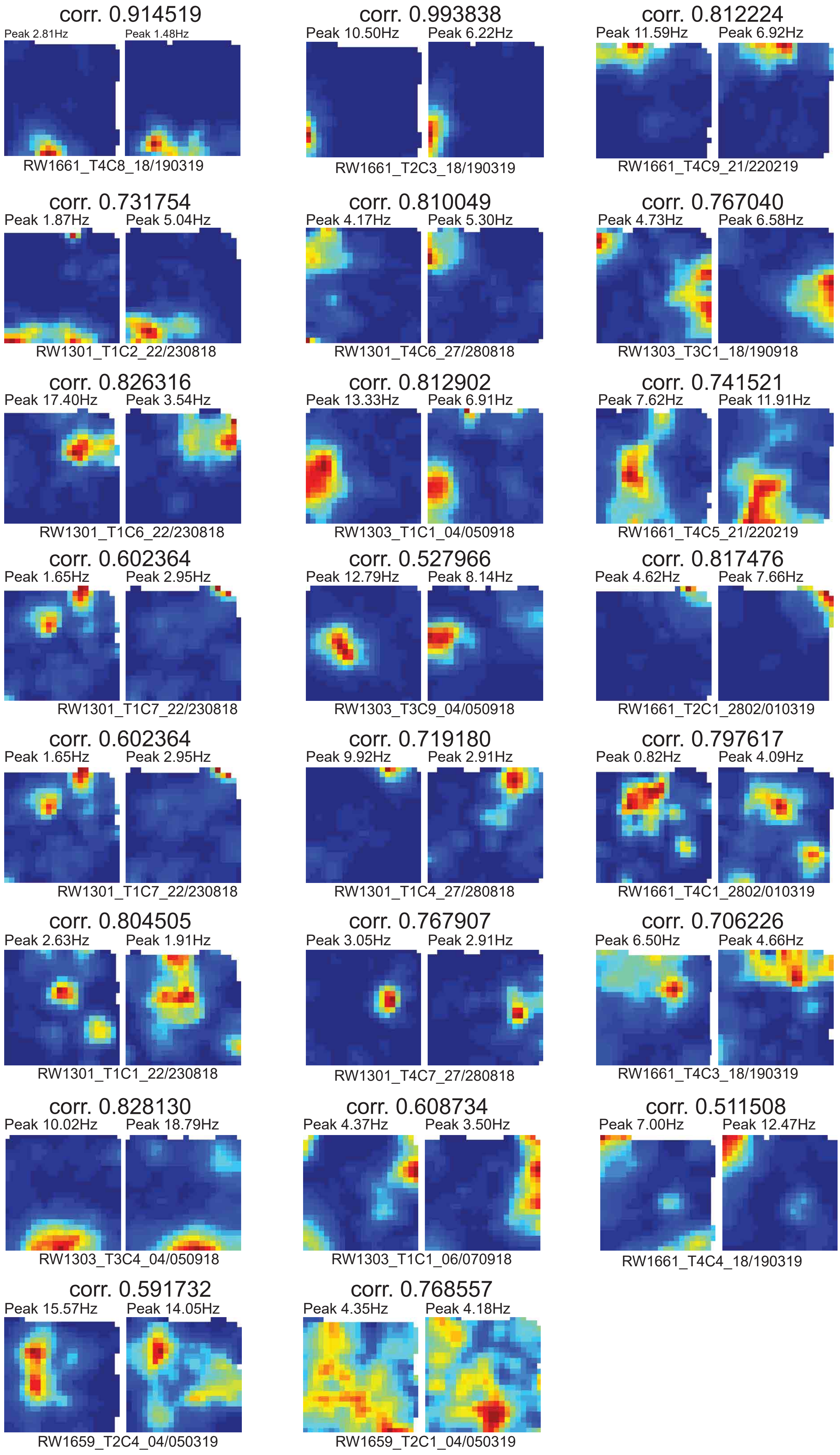
corr. 0.914519
Peak 2.81Hz
Peak 1.48Hz
RW1661_T4C8_18/190319
corr. 0.993838
Peak 10.50Hz
Peak 6.22Hz
RW1661_T2C3_18/190319
corr. 0.812224
Peak 11.59Hz
Peak 6.92Hz
RW1661_T4C9_21/220219
corr. 0.731754
Peak 1.87Hz
Peak 5.04Hz
RW1301_T1C2_22/230818
corr. 0.810049
Peak 4.17Hz
Peak 5.30Hz
RW1301_T4C6_27/280818
corr. 0.767040
Peak 4.73Hz
Peak 6.58Hz
RW1303_T3C1_18/190918
corr. 0.826316
Peak 17.40Hz
Peak 3.54Hz
RW1301_T1C6_22/230818
corr. 0.812902
Peak 13.33Hz
Peak 6.91Hz
RW1303_T1C1_04/050918
corr. 0.741521
Peak 7.62Hz
Peak 11.91Hz
RW1661_T4C5_21/220219
corr. 0.602364
Peak 1.65Hz
Peak 2.95Hz
RW1301_T1C7_22/230818
corr. 0.527966
Peak 12.79Hz
Peak 8.14Hz
RW1303_T3C9_04/050918
corr. 0.817476
Peak 4.62Hz
Peak 7.66Hz
RW1661_T2C1_2802/010319
corr. 0.602364
Peak 1.65Hz
Peak 2.95Hz
RW1301_T1C7_22/230818
corr. 0.719180
Peak 9.92Hz
Peak 2.91Hz
RW1301_T1C4_27/280818
corr. 0.797617
Peak 0.82Hz
Peak 4.09Hz
RW1661_T4C1_2802/010319
corr. 0.804505
Peak 2.63Hz
Peak 1.91Hz
RW1301_T1C1_22/230818
corr. 0.767907
Peak 3.05Hz
Peak 2.91Hz
RW1301_T4C7_27/280818
corr. 0.706226
Peak 6.50Hz
Peak 4.66Hz
RW1661_T4C3_18/190319
corr. 0.828130
Peak 10.02Hz
Peak 18.79Hz
RW1303_T3C4_04/050918
corr. 0.608734
Peak 4.37Hz
Peak 3.50Hz
RW1303_T1C1_06/070918
corr. 0.511508
Peak 7.00Hz
Peak 12.47Hz
RW1661_T4C4_18/190319
corr. 0.591732
Peak 15.57Hz
Peak 14.05Hz
RW1659_T2C4_04/050319
corr. 0.768557
Peak 4.35Hz
Peak 4.18Hz
RW1659_T2C1_04/050319

**Figure S3-4**

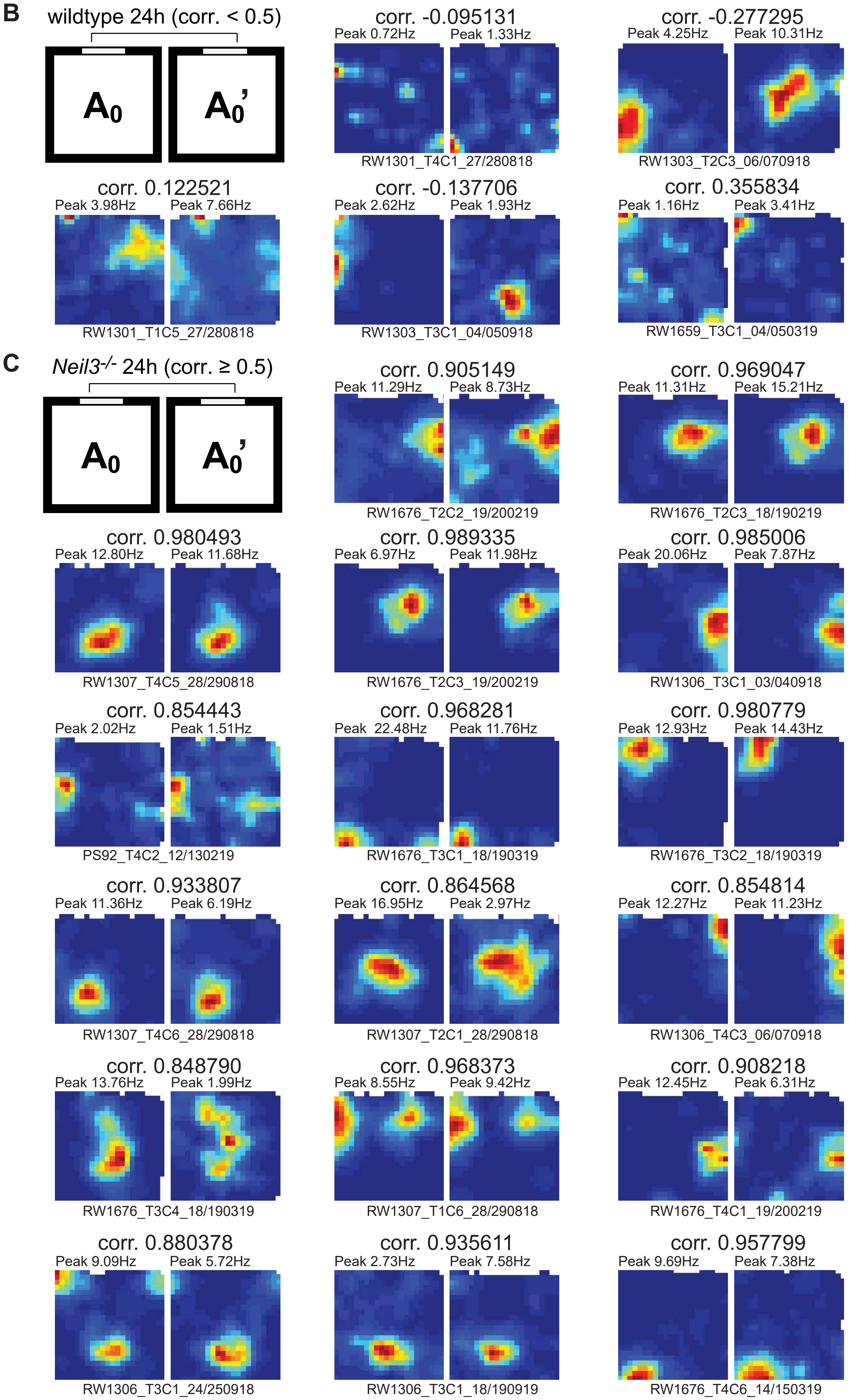
B
wildtype 24h (corr. < 0.5)
A0
A0’
corr. -0.095131
Peak 0.72Hz
Peak 1.33Hz
RW1301_T4C1_27/280818
corr. -0.277295
Peak 4.25Hz
Peak 10.31Hz
RW1303_T2C3_06/070918
corr. 0.122521
Peak 3.98Hz
Peak 7.66Hz
RW1301_T1C5_27/280818
corr. -0.137706
Peak 2.62Hz
Peak 1.93Hz
RW1303_T3C1_04/050918
corr. 0.355834
Peak 1.16Hz
Peak 3.41Hz
RW1659_T3C1_04/050319
C
Neil3-/- 24h (corr. ≥ 0.5)
A0
A0’
corr. 0.905149
Peak 11.29Hz
Peak 8.73Hz
RW1676_T2C2_19/200219
corr. 0.969047
Peak 11.31Hz
Peak 15.21Hz
RW1676_T2C3_18/190219
corr. 0.980493
Peak 12.80Hz
Peak 11.68Hz
RW1307_T4C5_28/290818
corr. 0.989335
Peak 6.97Hz
Peak 11.98Hz
RW1676_T2C3_19/200219
corr. 0.985006
Peak 20.06Hz
Peak 7.87Hz
RW1306_T3C1_03/040918
corr. 0.854443
Peak 2.02Hz
Peak 1.51Hz
PS92_T4C2_12/130219
corr. 0.968281
Peak 22.48Hz
Peak 11.76Hz
RW1676_T3C1_18/190319
corr. 0.980779
Peak 12.93Hz
Peak 14.43Hz
RW1676_T3C2_18/190319
corr. 0.933807
Peak 11.36Hz
Peak 6.19Hz
RW1307_T4C6_28/290818
corr. 0.864568
Peak 16.95Hz
Peak 2.97Hz
RW1307_T2C1_28/290818
corr. 0.854814
Peak 12.27Hz
Peak 11.23Hz
RW1306_T4C3_06/070918
corr. 0.848790
Peak 13.76Hz
Peak 1.99Hz
RW1676_T3C4_18/190319
corr. 0.968373
Peak 8.55Hz
Peak 9.42Hz
RW1307_T1C6_28/290818
corr. 0.908218
Peak 12.45Hz
Peak 6.31Hz
RW1676_T4C1_19/200219
corr. 0.880378
Peak 9.09Hz
Peak 5.72Hz
RW1306_T3C1_24/250918
corr. 0.935611
Peak 2.73Hz
Peak 7.58Hz
RW1306_T3C1_18/190919
corr. 0.957799
Peak 9.69Hz
Peak 7.38Hz
RW1676_T4C6_14/150319

**Figure S3-5**

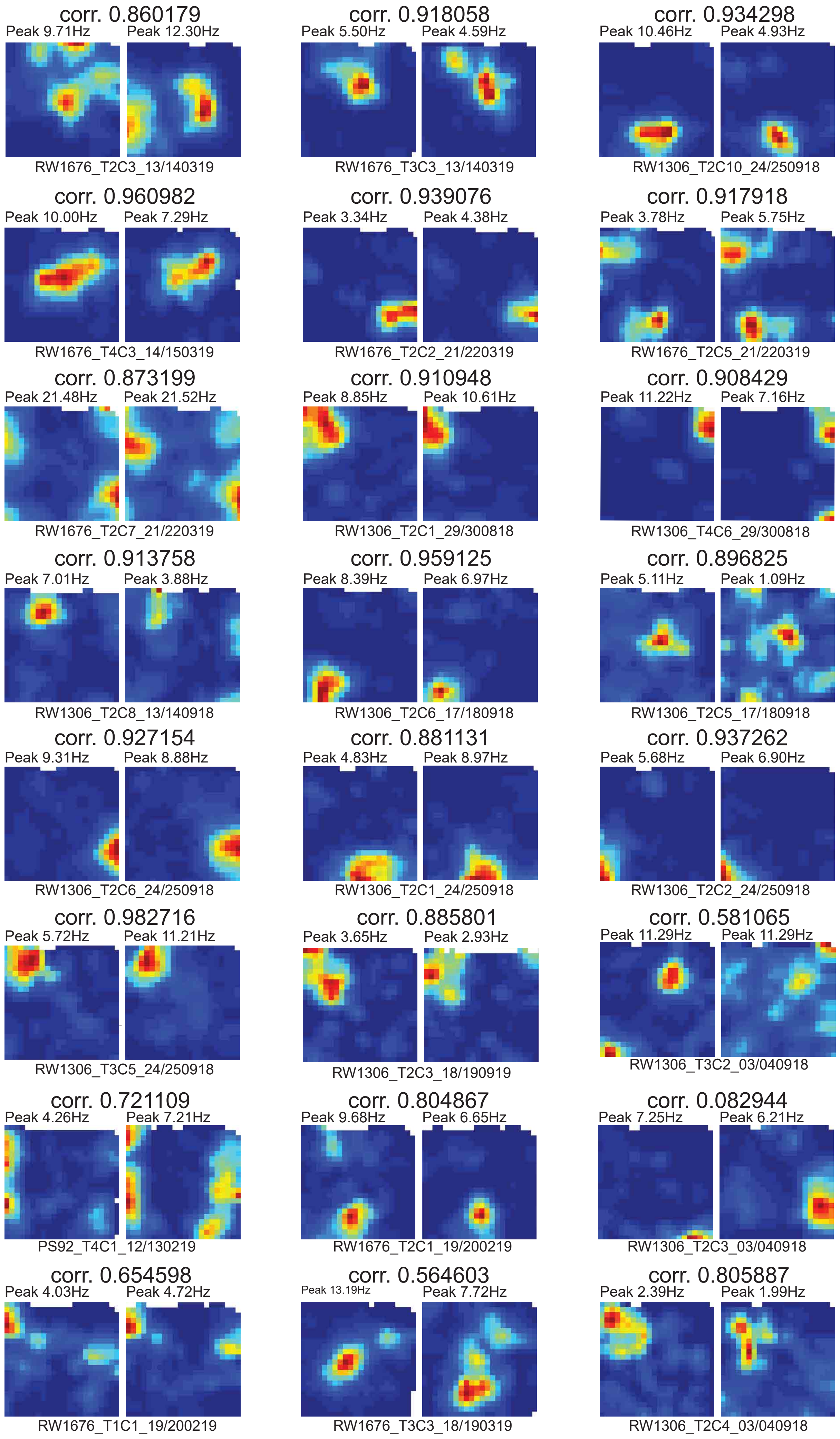
corr. 0.860179
Peak 9.71Hz
Peak 12.30Hz
RW1676_T2C3_13/140319
corr. 0.918058
Peak 5.50Hz
Peak 4.59Hz
RW1676_T3C3_13/140319
corr. 0.934298
Peak 10.46Hz
Peak 4.93Hz
RW1306_T2C10_24/250918
corr. 0.960982
Peak 10.00Hz
Peak 7.29Hz
RW1676_T4C3_14/150319
corr. 0.939076
Peak 3.34Hz
Peak 4.38Hz
RW1676_T2C2_21/220319
corr. 0.917918
Peak 3.78Hz
Peak 5.75Hz
RW1676_T2C5_21/220319
corr. 0.873199
Peak 21.48Hz
Peak 21.52Hz
RW1676_T2C7_21/220319
corr. 0.910948
Peak 8.85Hz
Peak 10.61Hz
RW1306_T2C1_29/300818
corr. 0.908429
Peak 11.22Hz
Peak 7.16Hz
RW1306_T4C6_29/300818
corr. 0.913758
Peak 7.01Hz
Peak 3.88Hz
RW1306_T2C8_13/140918
corr. 0.959125
Peak 8.39Hz
Peak 6.97Hz
RW1306_T2C6_17/180918
corr. 0.896825
Peak 5.11Hz
Peak 1.09Hz
RW1306_T2C5_17/180918
corr. 0.927154
Peak 9.31Hz
Peak 8.88Hz
RW1306_T2C6_24/250918
corr. 0.881131
Peak 4.83Hz
Peak 8.97Hz
RW1306_T2C1_24/250918
corr. 0.937262
Peak 5.68Hz
Peak 6.90Hz
RW1306_T2C2_24/250918
corr. 0.982716
Peak 5.72Hz
Peak 11.21Hz
RW1306_T3C5_24/250918
corr. 0.885801
Peak 3.65Hz
Peak 2.93Hz
RW1306_T2C3_18/190919
corr. 0.581065
Peak 11.29Hz
Peak 11.29Hz
RW1306_T3C2_03/040918
corr. 0.721109
Peak 4.26Hz
Peak 7.21Hz
PS92_T4C1_12/130219
corr. 0.804867
Peak 9.68Hz
Peak 6.65Hz
RW1676_T2C1_19/200219
corr. 0.082944
Peak 7.25Hz
Peak 6.21Hz
RW1306_T2C3_03/040918
corr. 0.654598
Peak 4.03Hz
Peak 4.72Hz
RW1676_T1C1_19/200219
corr. 0.564603
Peak 13.19Hz
Peak 7.72Hz
RW1676_T3C3_18/190319
corr. 0.805887
Peak 2.39Hz
Peak 1.99Hz
RW1306_T2C4_03/040918

**Figure S3-6**

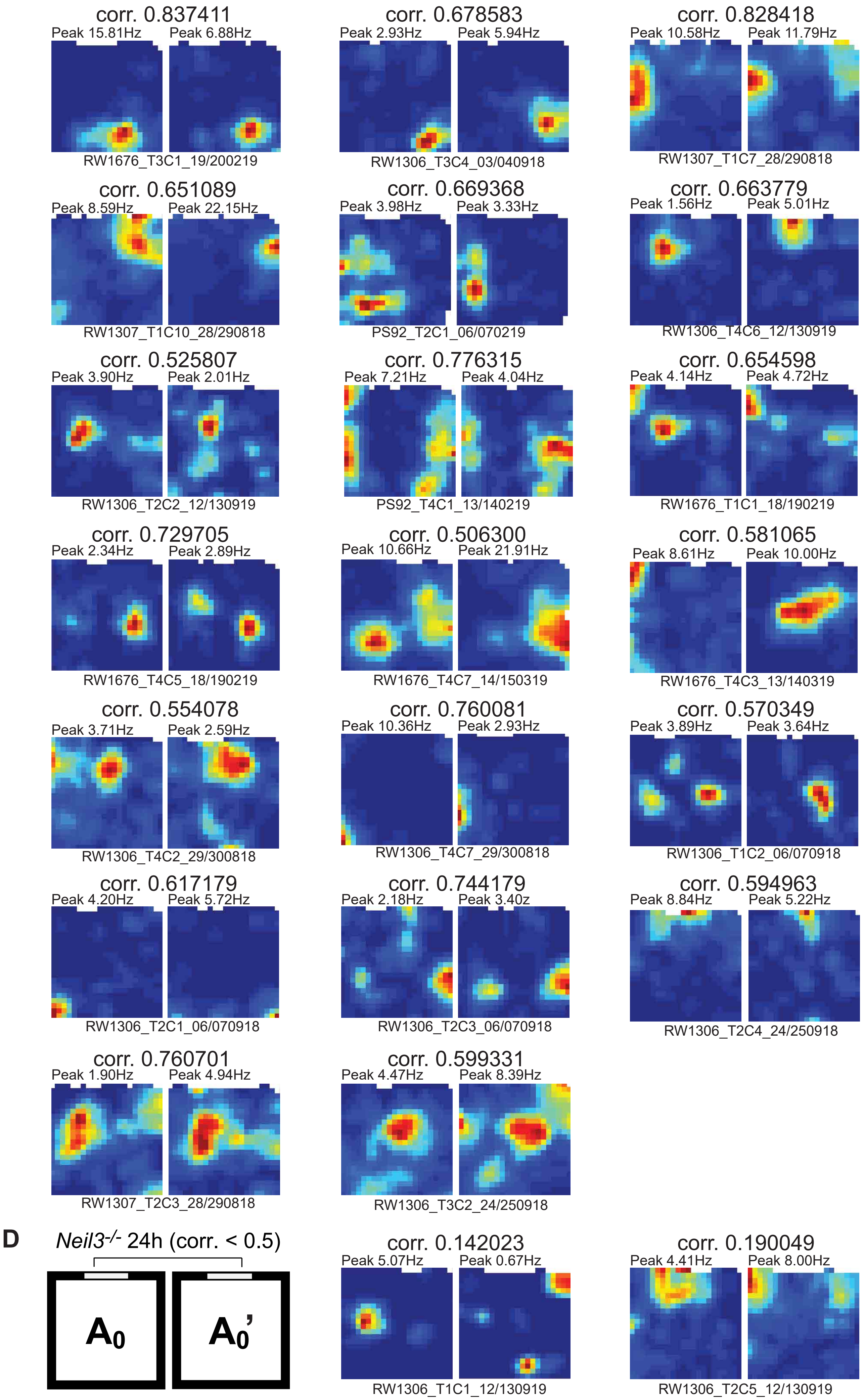
corr. 0.837411
Peak 15.81Hz
Peak 6.88Hz
RW1676_T3C1_19/200219
corr. 0.678583
Peak 2.93Hz
Peak 5.94Hz
RW1306_T3C4_03/040918
corr. 0.828418
Peak 10.58Hz
Peak 11.79Hz
RW1307_T1C7_28/290818
corr. 0.651089
Peak 8.59Hz
Peak 22.15Hz
RW1307_T1C10_28/290818
corr. 0.669368
Peak 3.98Hz
Peak 3.33Hz
PS92_T2C1_06/070219
corr. 0.663779
Peak 1.56Hz
Peak 5.01Hz
RW1306_T4C6_12/130919
corr. 0.525807
Peak 3.90Hz
Peak 2.01Hz
RW1306_T2C2_12/130919
corr. 0.776315
Peak 7.21Hz
Peak 4.04Hz
PS92_T4C1_13/140219
corr. 0.654598
Peak 4.14Hz
Peak 4.72Hz
RW1676_T1C1_18/190219
corr. 0.729705
Peak 2.34Hz
Peak 2.89Hz
RW1676_T4C5_18/190219
corr. 0.506300
Peak 10.66Hz
Peak 21.91Hz
RW1676_T4C7_14/150319
corr. 0.581065
Peak 8.61Hz
Peak 10.00Hz
RW1676_T4C3_13/140319
corr. 0.554078
Peak 3.71Hz
Peak 2.59Hz
RW1306_T4C2_29/300818
corr. 0.760081
Peak 10.36Hz
Peak 2.93Hz
RW1306_T4C7_29/300818
corr. 0.570349
Peak 3.89Hz
Peak 3.64Hz
RW1306_T1C2_06/070918
corr. 0.617179
Peak 4.20Hz
Peak 5.72Hz
RW1306_T2C1_06/070918
corr. 0.744179
Peak 2.18Hz
Peak 3.40z
RW1306_T2C3_06/070918
corr. 0.594963
Peak 8.84Hz
Peak 5.22Hz
RW1306_T2C4_24/250918
corr. 0.760701
Peak 1.90Hz
Peak 4.94Hz
RW1307_T2C3_28/290818
corr. 0.599331
Peak 4.47Hz
Peak 8.39Hz
RW1306_T3C2_24/250918
D
Neil3-/- 24h (corr. < 0.5)
A0
A0'
corr. 0.142023
Peak 5.07Hz
Peak 0.67Hz
RW1306_T1C1_12/130919
corr. 0.190049
Peak 4.41Hz
Peak 8.00Hz
RW1306_T2C5_12/130919

Figure S3-7

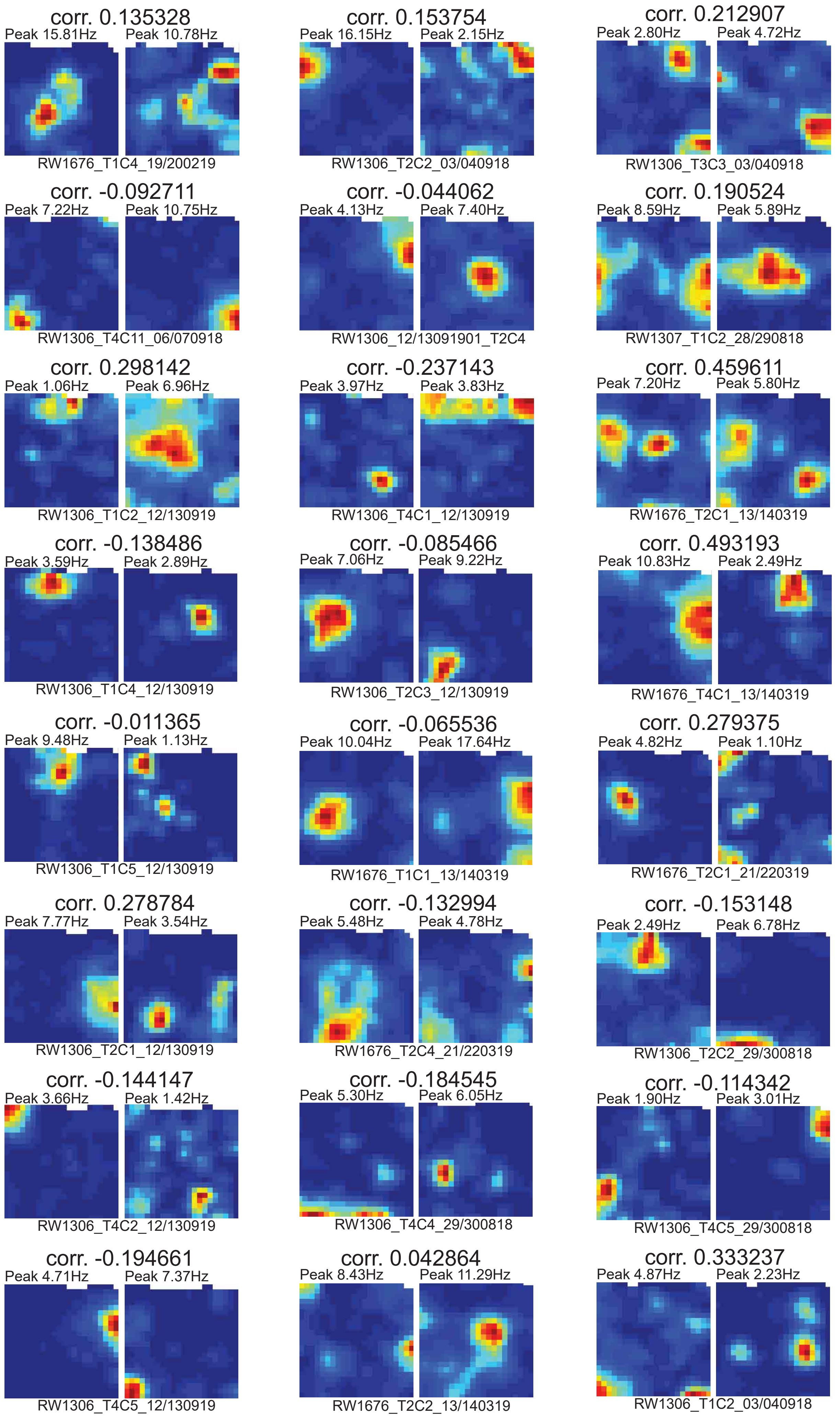

corr. 0.135328
Peak 15.81Hz
Peak 10.78Hz
RW1676_T1C4_19/200219
corr. 0.153754
Peak 16.15Hz
Peak 2.15Hz
RW1306_T2C2_03/040918
corr. 0.212907
Peak 2.80Hz
Peak 4.72Hz
RW1306_T3C3_03/040918
corr. -0.092711
Peak 7.22Hz
Peak 10.75Hz
RW1306_T4C11_06/070918
corr. -0.044062
Peak 4.13Hz
Peak 7.40Hz
RW1306_12/13091901_T2C4
corr. 0.190524
Peak 8.59Hz
Peak 5.89Hz
RW1307_T1C2_28/290818
corr. 0.298142
Peak 1.06Hz
Peak 6.96Hz
RW1306_T1C2_12/130919
corr. -0.237143
Peak 3.97Hz
Peak 3.83Hz
RW1306_T4C1_12/130919
corr. 0.459611
Peak 7.20Hz
Peak 5.80Hz
RW1676_T2C1_13/140319
corr. -0.138486
Peak 3.59Hz
Peak 2.89Hz
RW1306_T1C4_12/130919
corr. -0.085466
Peak 7.06Hz
Peak 9.22Hz
RW1306_T2C3_12/130919
corr. 0.493193
Peak 10.83Hz
Peak 2.49Hz
RW1676_T4C1_13/140319
corr. -0.011365
Peak 9.48Hz
Peak 1.13Hz
RW1306_T1C5_12/130919
corr. -0.065536
Peak 10.04Hz
Peak 17.64Hz
RW1676_T1C1_13/140319
corr. 0.279375
Peak 4.82Hz
Peak 1.10Hz
RW1676_T2C1_21/220319
corr. 0.278784
Peak 7.77Hz
Peak 3.54Hz
RW1306_T2C1_12/130919
corr. -0.132994
Peak 5.48Hz
Peak 4.78Hz
RW1676_T2C4_21/220319
corr. -0.153148
Peak 2.49Hz
Peak 6.78Hz
RW1306_T2C2_29/300818
corr. -0.144147
Peak 3.66Hz
Peak 1.42Hz
RW1306_T4C2_12/130919
corr. -0.184545
Peak 5.30Hz
Peak 6.05Hz
RW1306_T4C4_29/300818
corr. -0.114342
Peak 1.90Hz
Peak 3.01Hz
RW1306_T4C5_29/300818
corr. -0.194661
Peak 4.71Hz
Peak 7.37Hz
RW1306_T4C5_12/130919
corr. 0.042864
Peak 8.43Hz
Peak 11.29Hz
RW1676_T2C2_13/140319
corr. 0.333237
Peak 4.87Hz
Peak 2.23Hz
RW1306_T1C2_03/040918

**Figure S3-8**

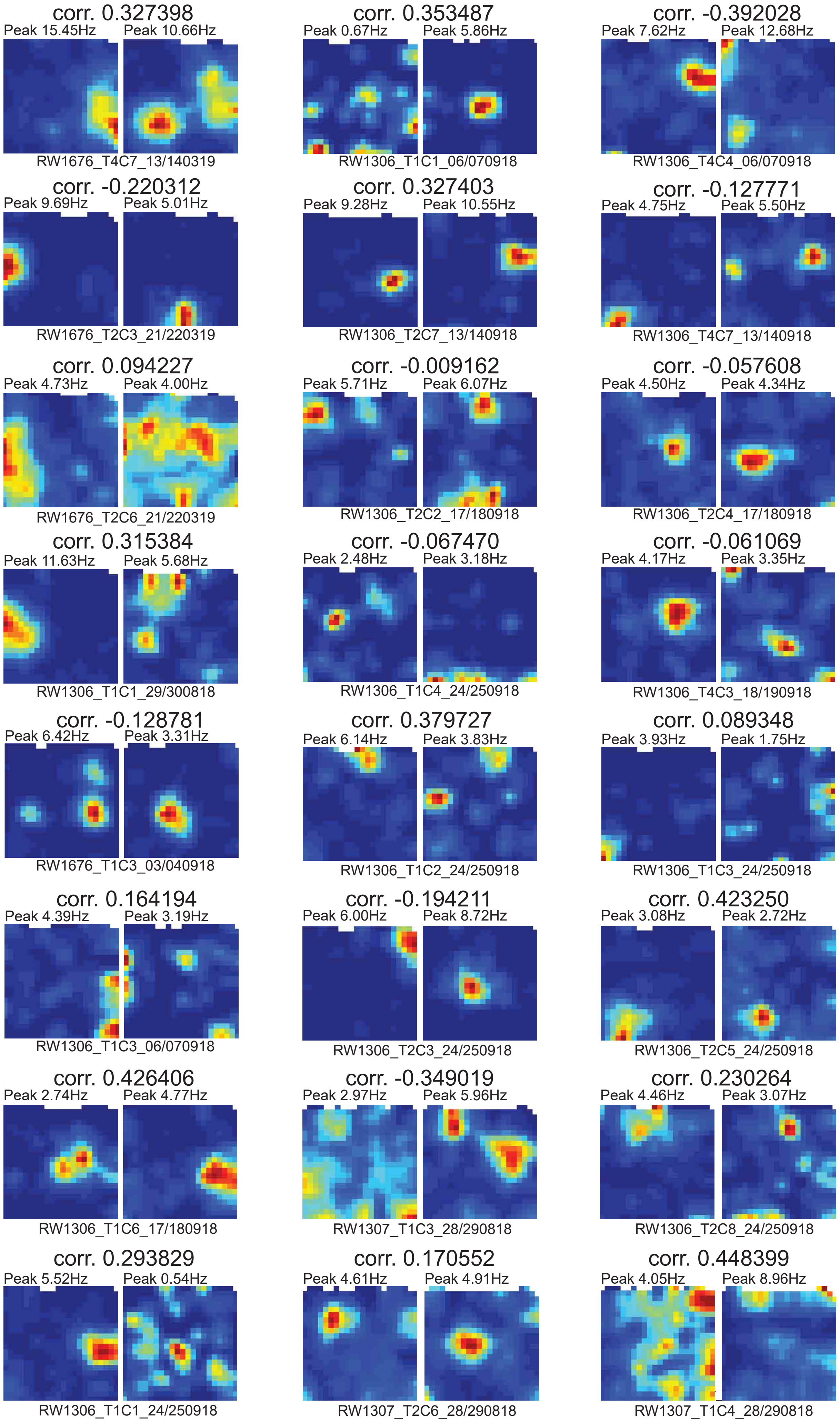
corr. 0.327398
Peak 15.45Hz
Peak 10.66Hz
RW1676_T4C7_13/140319
corr. 0.353487
Peak 0.67Hz
Peak 5.86Hz
RW1306_T1C1_06/070918
corr. -0.392028
Peak 7.62Hz
Peak 12.68Hz
RW1306_T4C4_06/070918
corr. -0.220312
Peak 9.69Hz
Peak 5.01Hz
RW1676_T2C3_21/220319
corr. 0.327403
Peak 9.28Hz
Peak 10.55Hz
RW1306_T2C7_13/140918
corr. -0.127771
Peak 4.75Hz
Peak 5.50Hz
RW1306_T4C7_13/140918
corr. 0.094227
Peak 4.73Hz
Peak 4.00Hz
RW1676_T2C6_21/220319
corr. -0.009162
Peak 5.71Hz
Peak 6.07Hz
RW1306_T2C2_17/180918
corr. -0.057608
Peak 4.50Hz
Peak 4.34Hz
RW1306_T2C4_17/180918
corr. 0.315384
Peak 11.63Hz
Peak 5.68Hz
RW1306_T1C1_29/300818
corr. -0.067470
Peak 2.48Hz
Peak 3.18Hz
RW1306_T1C4_24/250918
corr. -0.061069
Peak 4.17Hz
Peak 3.35Hz
RW1306_T4C3_18/190918
corr. -0.128781
Peak 6.42Hz
Peak 3.31Hz
RW1676_T1C3_03/040918
corr. 0.379727
Peak 6.14Hz
Peak 3.83Hz
RW1306_T1C2_24/250918
corr. 0.089348
Peak 3.93Hz
Peak 1.75Hz
RW1306_T1C3_24/250918
corr. 0.164194
Peak 4.39Hz
Peak 3.19Hz
RW1306_T1C3_06/070918
corr. -0.194211
Peak 6.00Hz
Peak 8.72Hz
RW1306_T2C3_24/250918
corr. 0.423250
Peak 3.08Hz
Peak 2.72Hz
RW1306_T2C5_24/250918
corr. 0.426406
Peak 2.74Hz
Peak 4.77Hz
RW1306_T1C6_17/180918
corr. -0.349019
Peak 2.97Hz
Peak 5.96Hz
RW1307_T1C3_28/290818
corr. 0.230264
Peak 4.46Hz
Peak 3.07Hz
RW1306_T2C8_24/250918
corr. 0.293829
Peak 5.52Hz
Peak 0.54Hz
RW1306_T1C1_24/250918
corr. 0.170552
Peak 4.61Hz
Peak 4.91Hz
RW1307_T2C6_28/290818
corr. 0.448399
Peak 4.05Hz
Peak 8.96Hz
RW1307_T1C4_28/290818

**Figure S3-9**

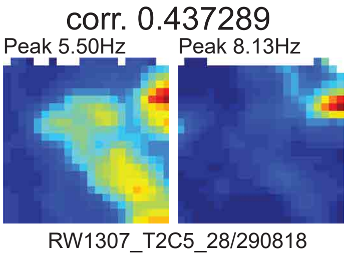


**Supplement Figure 3. 24h-comparison of place field stability in wildtype and *Neil3*$^{-/-}$ mice.** Rate maps of wildtype place cells in trial $A_0$ (Day1) and $A_0$' (Day2) with a spatial cross-correlation coefficient above and below 0.5 were listed in (**A**) and (**B**). Similarly, rate maps of *Neil3*$^{-/-}$ place cells in $A_0$ (Day1) and $A_0$' (Day2) with a spatial cross-correlation coefficient above and below 0.5 were listed in (**C**) and (**D**).

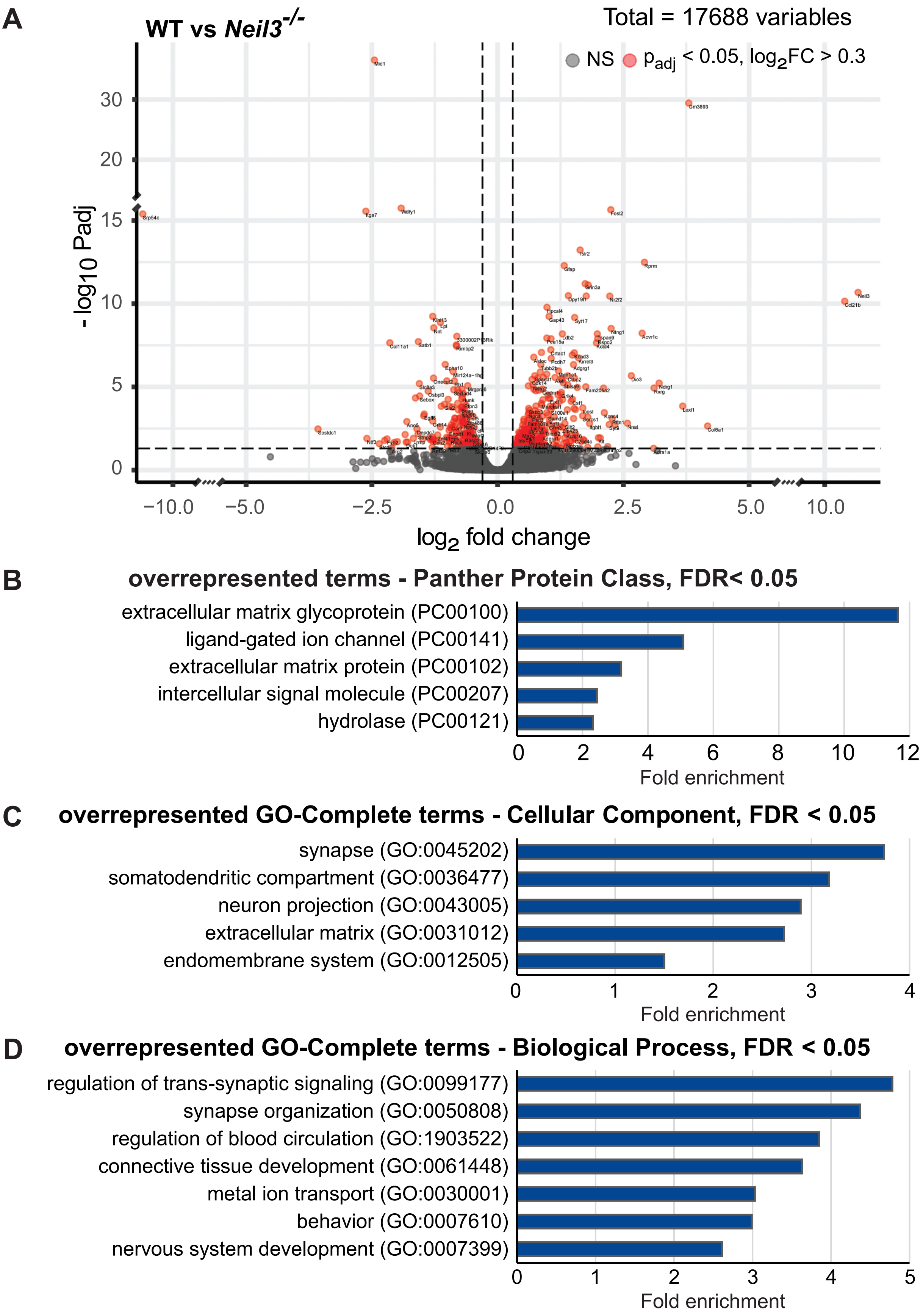


**Supplement Figure 4. Analysis of differentially expressed genes (DEGs) in "baseline" condition.** (**A**) Volcano plot showing differentially expressed genes (n = 2, $p_{adj} < 0.05$ / $log_2FC \geq 0.3$) as red dots. (**B**) Significantly enriched PANTHER Protein Class terms (Binomial, FDR < 0.05, fold enrichment ≥ 2, overrepresentation based on DEGs with $p_{adj} < 0.05$ / $log_2FC \geq 0.3$). (**C**) Significantly enriched Gene Ontology Complete Cellular Component ancestral terms (Binomial, Bonferroni, fold enrichment ≥ 2, overrepresentation based on DEGs with $p_{adj} < 0.05$ / $log_2FC \geq 0.3$). (**D**) Significantly enriched Gene Ontology Complete Biological Process ancestral terms (Binomial, Bonferroni, overrepresentation based on DEGs with $p_{adj} < 0.05$ / $log_2FC \geq 0.3$).

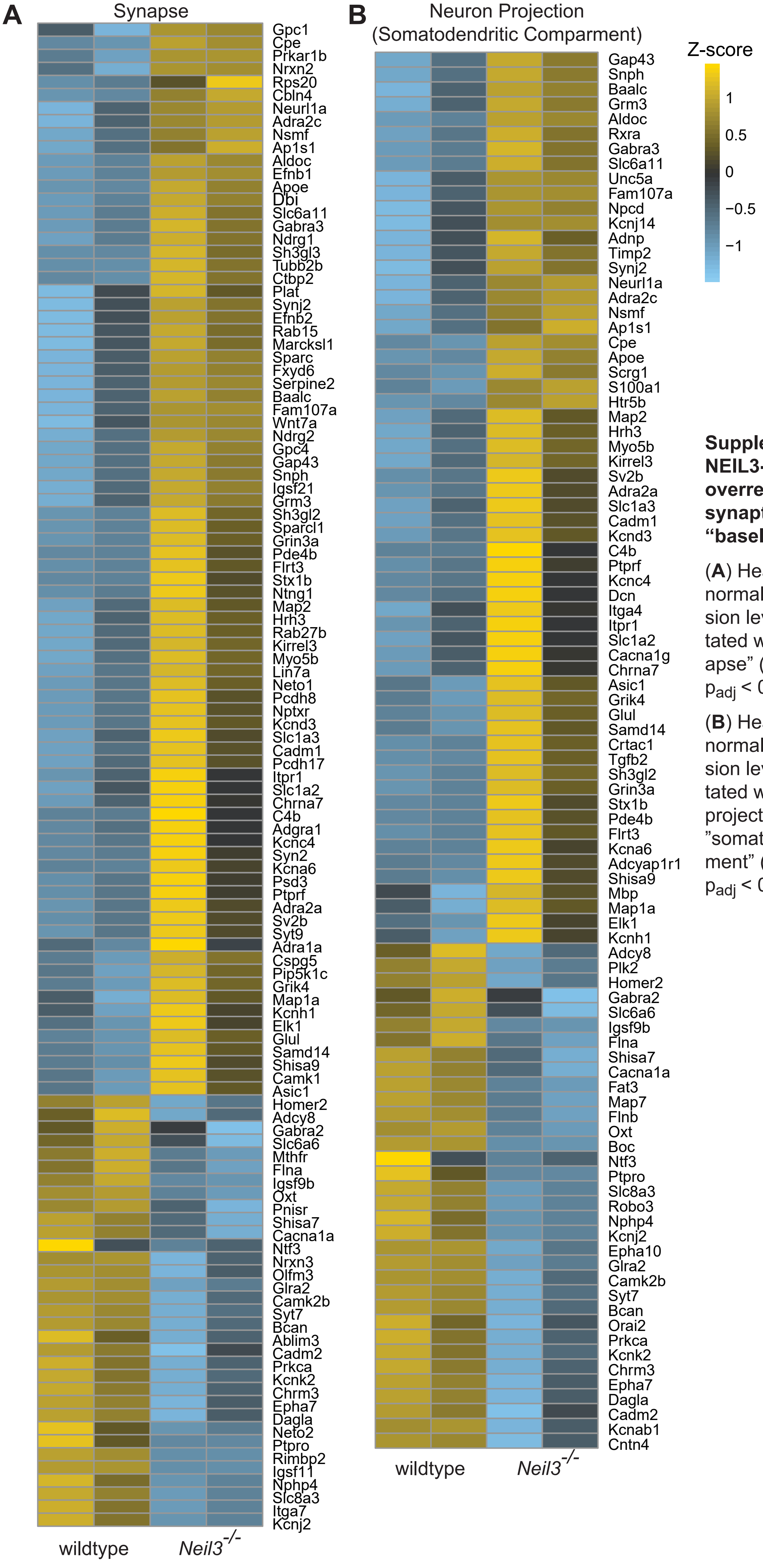


**Supplement Figure 5. NEIL3-dependent DEGs overrepresented in GO-CC synaptic components in "baseline" condition.**

(**A**) Heatmap of z-score normalized FPKM expression levels of genes annotated with the term "synapse" (GO:0045202, n=2, $p_{adj} < 0.05$, $\log_2 FC \geq 0.3$).

(**B**) Heatmap of z-score normalized FPKM expression levels of genes annotated with the term "neuron projection" (GO:0043005) / "somatodendritic compartment" (GO:0036477, n=2, $p_{adj} < 0.05$, $\log_2 FC \geq 0.3$).

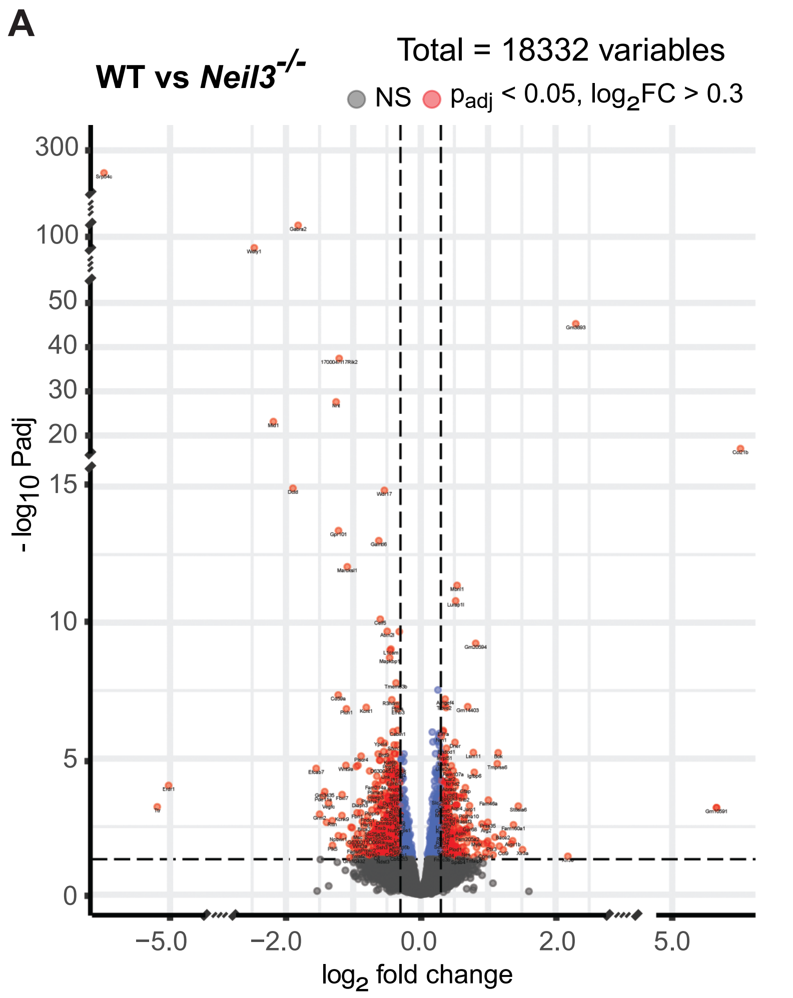


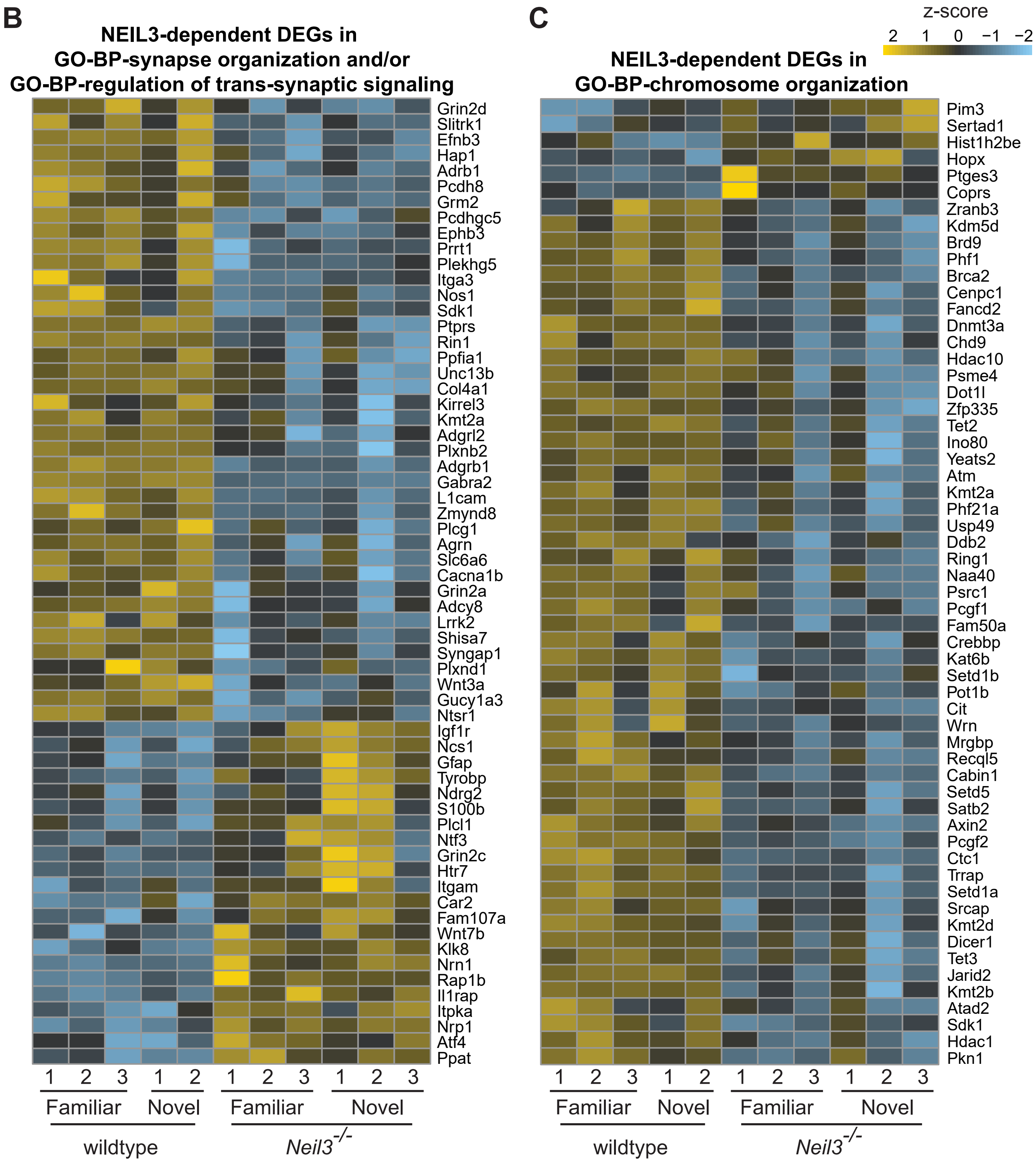


**Supplement Figure 6. Analysis of differentially expressed genes in wildtype and *Neil3*$^{-/-}$ mice after spatial exploration.** (**A**) Volcano plot showing differentially expressed genes (n=5 for wildtype, n=6 for *Neil3*$^{-/-}$, $p_{adj}<0.05$ / $log_2FC \geq 0.3$) as red dots. (**B**) Heatmap of z-score normalized FPKM expression levels of genes annotated with the GO-BP terms "synapse organization" (GO:0050808) and/or "regulation of trans-synaptic signaling" (GO:0099177, n=6 for wildtype, n=5 for *Neil3*$^{-/-}$, $p_{adj}$ < 0.05, $log_2FC \geq 0.3$). (**C**) Heatmap of z-score normalized FPKM expression levels of genes annotated with the GO-BP (Biological Process) term "chromosome organization" (GO:0051276, n=6 for wildtype, n=5 for *Neil3*$^{-/-}$, $p_{adj}$ < 0.05, $log_2FC \geq 0.3$). The conditions "familiar" and "novel" environment on the test day are subsumed due to virtually no difference in gene expression between both conditions within genotypes.

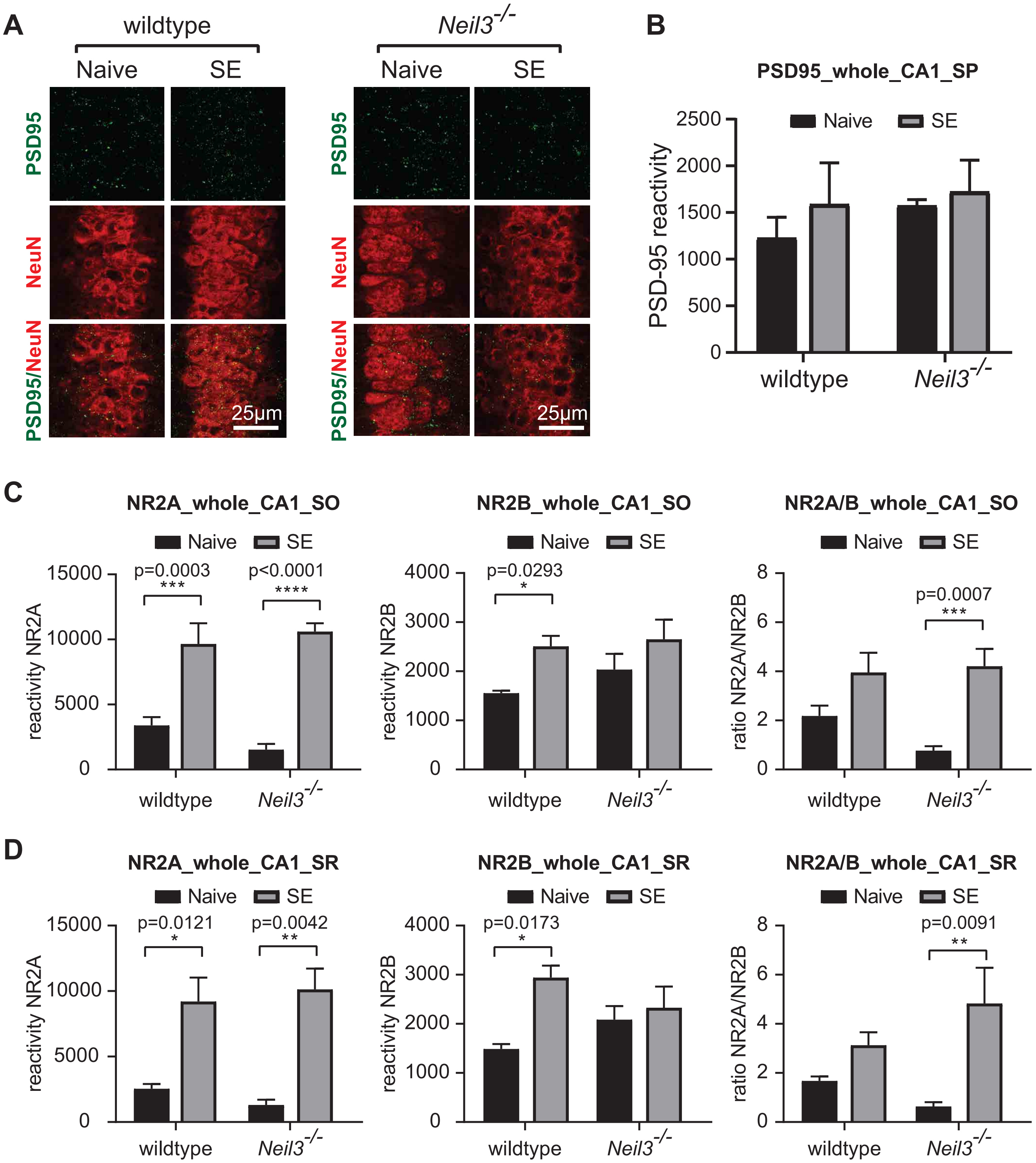


**Supplement Figure 7. Synaptic organization of PSD-95 and NMDA-receptor subunits in the hippocampal CA1 region of wildtype and *Neil3*$^{-/-}$ mice.** (**A**) Representative images showing dorsal hippocampal CA1 neurons that were immuno-stained with antibodies against PSD95 (green) and NeuN (red) from wt and *Neil3*$^{-/-}$ mice without (Naive) or after **s**patial **e**xploration (SE). (**B**) Bar diagrams showing the immunoreactivity (quantified in Imaris, see methods) of PSD95 at the pyramidal layer of CA1. (**C-D**) Bar diagrams showing the immunoreactivity (quantified in Imaris, see methods) of NR2A or NR2B at the stratum oriens (SO) and the stratum radiatum (SR) of CA1. The ratio of NR2A:NR2B was calculated for each animal individually. All statistics were conducted at animal level (3 animals with 2 hippocampal slices for each genotype, two-way-ANOVA, Sidak's correction, error bars indicate SEM).

### Supplement Table 1

| GO-Complete terms - Cellular Component (fold enrichment > 2) | Fold Enrichment | P-value |
|---|---|---|
| astrocyte projection (GO:0097449) | 14.56 | 4.25E-02 |
| integral component of postsynaptic membrane (GO:0099055) | 7.99 | 4.31E-14 |
| GABA-ergic synapse (GO:0098982) | 7.69 | 2.94E-07 |
| intrinsic component of postsynaptic membrane (GO:0098936) | 7.59 | 1.63E-13 |
| intrinsic component of presynaptic membrane (GO:0098889) | 7.28 | 1.95E-08 |
| intrinsic component of synaptic membrane (GO:0099240) | 7.06 | 9.10E-17 |
| integral component of synaptic membrane (GO:0099699) | 7.01 | 3.82E-15 |
| integral component of presynaptic membrane (GO:0099056) | 6.83 | 1.72E-06 |
| voltage-gated potassium channel complex (GO:0008076) | 6.65 | 1.86E-03 |
| integral component of postsynaptic specialization membrane (GO:0099060) | 6.65 | 2.26E-05 |
| integral component of postsynaptic density membrane (GO:0099061) | 6.56 | 6.35E-03 |
| presynaptic membrane (GO:0042734) | 6.38 | 1.08E-10 |
| intrinsic component of postsynaptic specialization membrane (GO:0098948) | 6.29 | 4.64E-05 |
| potassium channel complex (GO:0034705) | 6.21 | 1.26E-03 |
| neuron projection membrane (GO:0032589) | 6.16 | 3.09E-02 |
| intrinsic component of postsynaptic density membrane (GO:0099146) | 6.05 | 1.28E-02 |
| Schaffer collateral - CA1 synapse (GO:0098685) | 5.77 | 3.74E-04 |
| main axon (GO:0044304) | 5.48 | 2.95E-02 |
| postsynaptic specialization membrane (GO:0099634) | 5.48 | 1.06E-04 |
| postsynaptic density membrane (GO:0098839) | 5.28 | 1.61E-02 |
| synaptic membrane (GO:0097060) | 4.89 | 2.90E-16 |
| postsynaptic membrane (GO:0045211) | 4.8 | 1.95E-10 |
| cation channel complex (GO:0034703) | 4.7 | 6.12E-06 |
| glutamatergic synapse (GO:0098978) | 4.6 | 1.39E-15 |
| leading edge membrane (GO:0031256) | 4.38 | 8.96E-03 |
| presynapse (GO:0098793) | 4.21 | 4.93E-15 |
| neuron to neuron synapse (GO:0098984) | 4.19 | 4.03E-10 |
| ion channel complex (GO:0034702) | 4.15 | 3.14E-06 |
| postsynaptic specialization (GO:0099572) | 4.11 | 7.54E-10 |
| postsynaptic density (GO:0014069) | 4.04 | 2.03E-08 |
| asymmetric synapse (GO:0032279) | 4.01 | 2.48E-08 |
| transmembrane transporter complex (GO:1902495) | 3.93 | 9.19E-06 |
| postsynapse (GO:0098794) | 3.85 | 1.66E-15 |
| transporter complex (GO:1990351) | 3.84 | 1.44E-05 |
| synapse (GO:0045202) | 3.74 | 7.66E-32 |
| axon terminus (GO:0043679) | 3.74 | 2.66E-02 |
| neuronal cell body (GO:0043025) | 3.62 | 6.92E-13 |
| neuron projection terminus (GO:0044306) | 3.57 | 2.48E-02 |
| dendrite (GO:0030425) | 3.31 | 4.32E-10 |
| dendritic tree (GO:0097447) | 3.3 | 5.04E-10 |
| cell body (GO:0044297) | 3.26 | 2.61E-11 |
| distal axon (GO:0150034) | 3.24 | 3.92E-04 |
| axon (GO:0030424) | 3.22 | 2.10E-09 |
| somatodendritic compartment (GO:0036477) | 3.18 | 3.63E-14 |
| receptor complex (GO:0043235) | 3.05 | 1.18E-03 |
| plasma membrane protein complex (GO:0098797) | 3.02 | 1.78E-05 |
| collagen-containing extracellular matrix (GO:0062023) | 2.93 | 9.82E-03 |
| neuron projection (GO:0043005) | 2.89 | 2.47E-17 |
| extracellular matrix (GO:0031012) | 2.72 | 4.07E-03 |
| plasma membrane region (GO:0098590) | 2.67 | 7.00E-10 |
| secretory vesicle (GO:0099503) | 2.66 | 5.73E-04 |
| intrinsic component of plasma membrane (GO:0031226) | 2.63 | 1.10E-13 |
| cell junction (GO:0030054) | 2.56 | 9.61E-09 |
| integral component of plasma membrane (GO:0005887) | 2.55 | 1.29E-11 |
| plasma membrane bounded cell projection (GO:0120025) | 2.24 | 1.35E-12 |
| cell projection (GO:0042995) | 2.17 | 9.78E-13 |

### Supplement Table 2a

| GO-Complete terms - Biological Process (fold enrichment > 2.3) | Fold Enrichment | P-value |
|---|---|---|
| synaptic membrane adhesion (GO:0099560) | 13.8 | 1.60E-03 |
| proteoglycan metabolic process (GO:0006029) | 8.22 | 4.08E-04 |
| synapse assembly (GO:0007416) | 6.33 | 1.85E-02 |
| positive regulation of synaptic transmission (GO:0050806) | 5.92 | 2.09E-07 |
| potassium ion transmembrane transport (GO:0071805) | 5.52 | 5.88E-04 |
| regulation of synaptic vesicle exocytosis (GO:2000300) | 5.43 | 3.04E-02 |
| potassium ion transport (GO:0006813) | 5.21 | 2.14E-04 |
| regulation of neurotransmitter secretion (GO:0046928) | 5.18 | 3.37E-03 |
| cell-cell adhesion via plasma-membrane adhesion molecules (GO:0098742) | 5.06 | 5.69E-05 |
| regulation of muscle contraction (GO:0006937) | 4.81 | 3.63E-03 |
| modulation of chemical synaptic transmission (GO:0050804) | 4.79 | 5.43E-17 |
| regulation of synaptic vesicle cycle (GO:0098693) | 4.79 | 8.82E-03 |
| regulation of trans-synaptic signaling (GO:0099177) | 4.78 | 5.90E-17 |
| regulation of heart contraction (GO:0008016) | 4.71 | 9.35E-04 |
| neurotransmitter transport (GO:0006836) | 4.69 | 2.26E-03 |
| gliogenesis (GO:0042063) | 4.56 | 2.94E-05 |
| regulation of synaptic plasticity (GO:0048167) | 4.49 | 1.85E-04 |
| regulation of nervous system process (GO:0031644) | 4.47 | 4.25E-03 |
| memory (GO:0007613) | 4.47 | 4.39E-02 |
| regulation of neurotransmitter transport (GO:0051588) | 4.37 | 5.75E-03 |
| synapse organization (GO:0050808) | 4.37 | 3.17E-06 |
| regulation of endothelial cell migration (GO:0010594) | 4.37 | 2.66E-02 |
| regulation of axonogenesis (GO:0050770) | 4.28 | 1.80E-03 |
| axon guidance (GO:0007411) | 4.25 | 9.34E-04 |
| neuron projection guidance (GO:0097485) | 4.22 | 1.08E-03 |
| cartilage development (GO:0051216) | 4.19 | 2.13E-02 |
| trans-synaptic signaling (GO:0099537) | 4.15 | 1.22E-07 |
| learning or memory (GO:0007611) | 4.15 | 1.93E-05 |
| axon development (GO:0061564) | 4.14 | 1.31E-07 |
| glial cell differentiation (GO:0010001) | 4.03 | 3.44E-02 |
| cell projection morphogenesis (GO:0048858) | 3.98 | 1.47E-09 |
| regulation of regulated secretory pathway (GO:1903305) | 3.96 | 4.20E-02 |
| chemical synaptic transmission (GO:0007268) | 3.96 | 3.26E-06 |
| anterograde trans-synaptic signaling (GO:0098916) | 3.96 | 3.26E-06 |
| axonogenesis (GO:0007409) | 3.95 | 6.76E-06 |
| synaptic signaling (GO:0099536) | 3.93 | 2.57E-07 |
| cognition (GO:0050890) | 3.86 | 4.27E-05 |
| regulation of blood circulation (GO:1903522) | 3.85 | 4.39E-03 |
| plasma membrane bounded cell projection morphogenesis (GO:0120039) | 3.83 | 1.81E-08 |
| regulation of synapse organization (GO:0050807) | 3.78 | 1.60E-03 |
| cell part morphogenesis (GO:0032990) | 3.77 | 7.72E-09 |
| neuron projection morphogenesis (GO:0048812) | 3.77 | 5.47E-08 |
| cell morphogenesis involved in neuron differentiation (GO:0048667) | 3.77 | 4.09E-07 |
| calcium ion transport (GO:0006816) | 3.68 | 3.04E-02 |
| connective tissue development (GO:0061448) | 3.63 | 3.63E-02 |
| regulation of synapse structure or activity (GO:0050803) | 3.62 | 3.28E-03 |
| cell-cell adhesion (GO:0098609) | 3.59 | 2.96E-05 |
| regulation of exocytosis (GO:0017157) | 3.57 | 2.51E-02 |
| inorganic ion transmembrane transport (GO:0098660) | 3.54 | 3.69E-06 |
| inorganic cation transmembrane transport (GO:0098662) | 3.53 | 2.41E-05 |
| cell morphogenesis involved in differentiation (GO:0000904) | 3.52 | 1.98E-08 |
| regulation of neurotransmitter levels (GO:0001505) | 3.51 | 1.65E-04 |
| cell morphogenesis (GO:0000902) | 3.5 | 1.68E-11 |

### Supplement Table 2b

| | | |
|---|---|---|
| regulation of cell morphogenesis involved in differentiation (GO:0010769) | 3.48 | 6.16E-04 |
| negative regulation of neuron differentiation (GO:0045665) | 3.44 | 4.14E-02 |
| divalent metal ion transport (GO:0070838) | 3.41 | 2.67E-02 |
| divalent inorganic cation transport (GO:0072511) | 3.38 | 3.14E-02 |
| neuron projection development (GO:0031175) | 3.32 | 4.44E-09 |
| cation transmembrane transport (GO:0098655) | 3.29 | 1.20E-04 |
| monovalent inorganic cation transport (GO:0015672) | 3.21 | 2.26E-02 |
| negative regulation of neurogenesis (GO:0050768) | 3.2 | 1.44E-02 |
| regulation of system process (GO:0044057) | 3.18 | 1.34E-06 |
| cellular component morphogenesis (GO:0032989) | 3.11 | 1.20E-09 |
| negative regulation of nervous system development (GO:0051961) | 3.11 | 1.42E-02 |
| cell-cell signaling (GO:0007267) | 3.07 | 8.77E-09 |
| regulation of cell morphogenesis (GO:0022604) | 3.04 | 1.00E-04 |
| metal ion transport (GO:0030001) | 3.03 | 3.82E-05 |
| neuron development (GO:0048666) | 3 | 1.30E-08 |
| behavior (GO:0007610) | 2.99 | 1.88E-06 |
| negative regulation of cell development (GO:0010721) | 2.92 | 4.00E-02 |
| regulation of ion transport (GO:0043269) | 2.91 | 1.70E-06 |
| regulation of neuron differentiation (GO:0045664) | 2.88 | 9.39E-07 |
| regulation of ion transmembrane transport (GO:0034765) | 2.88 | 3.52E-03 |
| regulation of anatomical structure morphogenesis (GO:0022603) | 2.87 | 1.58E-10 |
| regulation of membrane potential (GO:0042391) | 2.85 | 1.56E-02 |
| ion transmembrane transport (GO:0034220) | 2.85 | 3.03E-04 |
| chemotaxis (GO:0006935) | 2.84 | 7.13E-03 |
| taxis (GO:0042330) | 2.82 | 8.06E-03 |
| developmental growth (GO:0048589) | 2.81 | 2.00E-02 |
| neuron differentiation (GO:0030182) | 2.79 | 5.03E-09 |
| regulation of nervous system development (GO:0051960) | 2.79 | 2.08E-09 |
| positive regulation of nervous system development (GO:0051962) | 2.78 | 1.46E-04 |
| positive regulation of neurogenesis (GO:0050769) | 2.72 | 2.15E-03 |
| growth (GO:0040007) | 2.72 | 3.78E-02 |
| cell adhesion (GO:0007155) | 2.72 | 1.49E-05 |
| neurogenesis (GO:0022008) | 2.72 | 1.35E-16 |
| calcium ion homeostasis (GO:0055074) | 2.71 | 2.60E-02 |
| plasma membrane bounded cell projection organization (GO:0120036) | 2.71 | 2.91E-08 |
| cellular calcium ion homeostasis (GO:0006874) | 2.7 | 4.08E-02 |
| cell projection organization (GO:0030030) | 2.7 | 9.45E-09 |
| regulation of neurogenesis (GO:0050767) | 2.69 | 4.62E-07 |
| cellular divalent inorganic cation homeostasis (GO:0072503) | 2.68 | 3.15E-02 |
| biological adhesion (GO:0022610) | 2.68 | 2.18E-05 |
| regulation of neuron projection development (GO:0010975) | 2.68 | 1.92E-03 |
| generation of neurons (GO:0048699) | 2.67 | 1.88E-14 |
| divalent inorganic cation homeostasis (GO:0072507) | 2.64 | 2.81E-02 |
| nervous system development (GO:0007399) | 2.61 | 6.10E-20 |
| regulation of transmembrane transport (GO:0034762) | 2.57 | 1.50E-02 |
| regulation of vesicle-mediated transport (GO:0060627) | 2.57 | 1.07E-02 |
| regulation of cell development (GO:0060284) | 2.56 | 3.92E-07 |
| positive regulation of cell development (GO:0010720) | 2.52 | 5.39E-03 |
| cation transport (GO:0006812) | 2.48 | 2.77E-03 |
| negative regulation of multicellular organismal process (GO:0051241) | 2.46 | 6.69E-08 |
| cell migration (GO:0016477) | 2.45 | 6.76E-04 |
| locomotion (GO:0040011) | 2.44 | 1.87E-06 |
| movement of cell or subcellular component (GO:0006928) | 2.34 | 6.46E-07 |
| localization of cell (GO:0051674) | 2.33 | 6.92E-04 |
| cell motility (GO:0048870) | 2.33 | 6.92E-04 |
| central nervous system development (GO:0007417) | 2.3 | 8.70E-03 |

### Supplement Table 3

| GO-Complete terms - Cellular Component (fold enrichment > 1.5) | Fold Enrichment | P-value |
|---|---|---|
| astrocyte projection (GO:0097449) | 6.49 | 4.12E-02 |
| cytosolic small ribosomal subunit (GO:0022627) | 5.08 | 7.24E-03 |
| cytosolic ribosome (GO:0022626) | 4.03 | 9.17E-05 |
| cytosolic large ribosomal subunit (GO:0022625) | 3.99 | 1.42E-02 |
| small ribosomal subunit (GO:0015935) | 3.33 | 3.84E-02 |
| integral component of postsynaptic membrane (GO:0099055) | 2.76 | 8.60E-03 |
| intrinsic component of postsynaptic membrane (GO:0098936) | 2.62 | 1.42E-02 |
| ribosomal subunit (GO:0044391) | 2.62 | 7.83E-03 |
| integral component of synaptic membrane (GO:0099699) | 2.53 | 7.37E-03 |
| intrinsic component of synaptic membrane (GO:0099240) | 2.45 | 7.96E-03 |
| ribosome (GO:0005840) | 2.35 | 1.83E-02 |
| postsynaptic membrane (GO:0045211) | 2.04 | 2.69E-02 |
| postsynaptic density (GO:0014069) | 1.96 | 2.40E-02 |
| glutamatergic synapse (GO:0098978) | 1.94 | 7.74E-03 |
| synaptic membrane (GO:0097060) | 1.93 | 1.28E-02 |
| asymmetric synapse (GO:0032279) | 1.93 | 2.73E-02 |
| neuron to neuron synapse (GO:0098984) | 1.93 | 2.10E-02 |
| postsynaptic specialization (GO:0099572) | 1.9 | 2.52E-02 |
| postsynapse (GO:0098794) | 1.88 | 1.09E-03 |
| synapse (GO:0045202) | 1.79 | 3.68E-06 |
| ribonucleoprotein complex (GO:1990904) | 1.78 | 7.96E-03 |
| axon (GO:0030424) | 1.77 | 7.84E-03 |
| neuron projection (GO:0043005) | 1.75 | 4.74E-06 |
| neuronal cell body (GO:0043025) | 1.74 | 1.24E-02 |
| somatodendritic compartment (GO:0036477) | 1.71 | 1.45E-03 |
| cell body (GO:0044297) | 1.66 | 1.65E-02 |
| dendrite (GO:0030425) | 1.65 | 3.43E-02 |
| dendritic tree (GO:0097447) | 1.65 | 3.65E-02 |
| cell junction (GO:0030054) | 1.64 | 2.82E-06 |
| cell projection (GO:0042995) | 1.54 | 8.05E-06 |
| plasma membrane bounded cell projection (GO:0120025) | 1.53 | 3.84E-05 |

### Supplement Table 4a

| GO-Complete terms - Biological Process (fold enrichment > 1.5) | Fold Enrichment | P-value |
|---|---|---|
| histone H3-K4 dimethylation (GO:0044648) | 20.76 | 6.23E-03 |
| histone H3-K4 monomethylation (GO:0097692) | 14.83 | 1.74E-02 |
| AV node cell to bundle of His cell signaling (GO:0086027) | 14.83 | 1.72E-02 |
| AV node cell action potential (GO:0086016) | 14.83 | 1.71E-02 |
| membrane depolarization during cardiac muscle cell action potential (GO:0086012) | 10.81 | 1.30E-02 |
| histone H3-K4 trimethylation (GO:0080182) | 9.56 | 1.83E-03 |
| peptidyl-lysine trimethylation (GO:0018023) | 6.31 | 2.77E-03 |
| histone H3-K4 methylation (GO:0051568) | 6.29 | 6.71E-03 |
| ribosomal large subunit assembly (GO:0000027) | 6.05 | 1.86E-02 |
| negative regulation of axon extension involved in axon guidance (GO:0048843) | 5.77 | 4.92E-02 |
| heterophilic cell-cell adhesion via plasma membrane cell adhesion molecules (GO:0007157) | 4.77 | 1.59E-02 |
| regulation of axon guidance (GO:1902667) | 4.51 | 3.73E-02 |
| negative regulation of axon extension (GO:0030517) | 4.42 | 4.23E-02 |
| cellular response to hydrogen peroxide (GO:0070301) | 4.32 | 1.54E-02 |
| histone lysine methylation (GO:0034968) | 3.99 | 2.38E-02 |
| ribosome assembly (GO:0042255) | 3.93 | 2.55E-02 |
| negative regulation of axonogenesis (GO:0050771) | 3.82 | 3.08E-02 |
| positive regulation of calcium ion transmembrane transport (GO:1904427) | 3.8 | 1.24E-02 |
| circadian regulation of gene expression (GO:0032922) | 3.71 | 3.64E-02 |
| peptidyl-lysine methylation (GO:0018022) | 3.61 | 2.63E-02 |
| histone methylation (GO:0016571) | 3.58 | 1.81E-02 |
| regulation of synapse assembly (GO:0051963) | 3.35 | 5.12E-03 |
| cellular response to reactive oxygen species (GO:0034614) | 2.96 | 4.43E-02 |
| positive regulation of transporter activity (GO:0032411) | 2.95 | 3.15E-02 |
| positive regulation of cation transmembrane transport (GO:1904064) | 2.92 | 6.10E-03 |
| negative regulation of neuron projection development (GO:0010977) | 2.82 | 8.92E-03 |
| negative regulation of developmental growth (GO:0048640) | 2.82 | 4.47E-02 |
| neuron projection guidance (GO:0097485) | 2.67 | 3.12E-03 |
| positive regulation of ion transmembrane transport (GO:0034767) | 2.65 | 1.65E-02 |
| negative regulation of cell projection organization (GO:0031345) | 2.65 | 9.12E-03 |
| synapse organization (GO:0050808) | 2.6 | 7.13E-04 |
| cell-cell adhesion via plasma-membrane adhesion molecules (GO:0098742) | 2.59 | 1.94E-02 |
| ribonucleoprotein complex assembly (GO:0022618) | 2.59 | 3.40E-02 |
| axon guidance (GO:0007411) | 2.57 | 6.68E-03 |
| peptidyl-lysine modification (GO:0018205) | 2.54 | 4.26E-03 |
| regulation of cell junction assembly (GO:1901888) | 2.53 | 1.47E-02 |
| ribonucleoprotein complex subunit organization (GO:0071826) | 2.49 | 4.75E-02 |
| cellular response to oxidative stress (GO:0034599) | 2.48 | 2.26E-02 |
| positive regulation of transmembrane transport (GO:0034764) | 2.47 | 1.15E-02 |
| regulation of synaptic plasticity (GO:0048167) | 2.45 | 1.93E-02 |
| covalent chromatin modification (GO:0016569) | 2.45 | 1.07E-03 |
| negative regulation of neuron differentiation (GO:0045665) | 2.42 | 1.07E-02 |
| histone modification (GO:0016570) | 2.41 | 1.74E-03 |
| neuron projection morphogenesis (GO:0048812) | 2.37 | 7.38E-05 |
| cell projection morphogenesis (GO:0048858) | 2.37 | 5.63E-05 |
| regulation of axonogenesis (GO:0050770) | 2.37 | 4.44E-02 |
| axonogenesis (GO:0007409) | 2.36 | 1.85E-03 |
| plasma membrane bounded cell projection morphogenesis (GO:0120039) | 2.35 | 9.53E-05 |
| cellular response to chemical stress (GO:0062197) | 2.34 | 2.45E-02 |
| regulation of neuron apoptotic process (GO:0043523) | 2.32 | 1.70E-02 |
| axon development (GO:0061564) | 2.3 | 1.69E-03 |
| regulation of neuron projection development (GO:0010975) | 2.27 | 1.63E-05 |
| cell part morphogenesis (GO:0032990) | 2.25 | 1.86E-04 |
| regulation of synapse structure or activity (GO:0050803) | 2.25 | 1.89E-02 |
| locomotory behavior (GO:0007626) | 2.25 | 4.47E-02 |

### Supplement Table 4b

| | | |
|---|---|---|
| regulation of transporter activity (GO:0032409) | 2.23 | 2.54E-02 |
| translation (GO:0006412) | 2.22 | 1.82E-02 |
| regulation of ion transmembrane transporter activity (GO:0032412) | 2.21 | 4.37E-02 |
| negative regulation of growth (GO:0045926) | 2.18 | 4.96E-02 |
| negative regulation of neurogenesis (GO:0050768) | 2.17 | 1.65E-02 |
| cell morphogenesis involved in neuron differentiation (GO:0048667) | 2.16 | 2.50E-03 |
| regulation of synapse organization (GO:0050807) | 2.16 | 4.43E-02 |
| negative regulation of nervous system development (GO:0051961) | 2.14 | 1.28E-02 |
| regulation of neuron differentiation (GO:0045664) | 2.12 | 1.22E-05 |
| positive regulation of neuron projection development (GO:0010976) | 2.11 | 1.60E-02 |
| modulation of chemical synaptic transmission (GO:0050804) | 2.11 | 1.23E-03 |
| regulation of trans-synaptic signaling (GO:0099177) | 2.1 | 1.28E-03 |
| regulation of nervous system development (GO:0051960) | 2.1 | 1.90E-07 |
| cellular component morphogenesis (GO:0032989) | 2.1 | 3.90E-04 |
| camera-type eye development (GO:0043010) | 2.09 | 2.92E-02 |
| peptide biosynthetic process (GO:0043043) | 2.09 | 3.61E-02 |
| positive regulation of nervous system development (GO:0051962) | 2.09 | 1.92E-04 |
| regulation of cation transmembrane transport (GO:1904062) | 2.08 | 2.14E-02 |
| regulation of neurogenesis (GO:0050767) | 2.08 | 1.67E-06 |
| neuron projection development (GO:0031175) | 2.06 | 1.46E-04 |
| peptide metabolic process (GO:0006518) | 2.05 | 9.47E-03 |
| response to oxidative stress (GO:0006979) | 2.03 | 4.19E-02 |
| negative regulation of cell development (GO:0010721) | 2.03 | 2.47E-02 |
| positive regulation of ion transport (GO:0043270) | 2.03 | 4.94E-02 |
| plasma membrane bounded cell projection organization (GO:0120036) | 2 | 1.69E-06 |
| brain development (GO:0007420) | 2 | 1.14E-03 |
| positive regulation of neurogenesis (GO:0050769) | 2 | 2.30E-03 |
| positive regulation of neuron differentiation (GO:0045666) | 1.99 | 1.23E-02 |
| trans-synaptic signaling (GO:0099537) | 1.99 | 4.40E-02 |
| head development (GO:0060322) | 1.99 | 6.15E-04 |
| regulation of cell morphogenesis (GO:0022604) | 1.98 | 5.23E-03 |
| regulation of cell development (GO:0060284) | 1.97 | 2.67E-06 |
| amide biosynthetic process (GO:0043604) | 1.97 | 2.51E-02 |
| visual system development (GO:0150063) | 1.97 | 4.23E-02 |
| sensory system development (GO:0048880) | 1.95 | 4.74E-02 |
| regulation of metal ion transport (GO:0010959) | 1.94 | 3.05E-02 |
| behavior (GO:0007610) | 1.94 | 1.45E-03 |
| cell-cell adhesion (GO:0098609) | 1.94 | 4.41E-02 |
| negative regulation of cell differentiation (GO:0045596) | 1.94 | 6.00E-04 |
| regulation of plasma membrane bounded cell projection organization (GO:0120035) | 1.94 | 4.21E-04 |
| cell projection organization (GO:0030030) | 1.93 | 5.40E-06 |
| neuron differentiation (GO:0030182) | 1.92 | 1.59E-05 |
| neuron development (GO:0048666) | 1.92 | 2.18E-04 |
| cell junction organization (GO:0034330) | 1.92 | 1.77E-02 |
| regulation of cell projection organization (GO:0031344) | 1.91 | 5.76E-04 |
| positive regulation of cell projection organization (GO:0031346) | 1.91 | 2.44E-02 |
| negative regulation of developmental process (GO:0051093) | 1.91 | 2.76E-05 |
| cell morphogenesis (GO:0000902) | 1.9 | 1.47E-03 |
| positive regulation of cell development (GO:0010720) | 1.9 | 3.15E-03 |
| chromatin organization (GO:0006325) | 1.88 | 5.65E-03 |
| cell morphogenesis involved in differentiation (GO:0000904) | 1.86 | 1.65E-02 |
| negative regulation of multicellular organismal process (GO:0051241) | 1.81 | 1.80E-05 |
| intracellular signal transduction (GO:0035556) | 1.8 | 1.79E-05 |
| regulation of transmembrane transport (GO:0034762) | 1.78 | 3.32E-02 |
| generation of neurons (GO:0048699) | 1.78 | 1.24E-06 |
| regulation of system process (GO:0044057) | 1.78 | 2.76E-02 |
| central nervous system development (GO:0007417) | 1.75 | 5.12E-03 |

**Supplement Table 4c**

| | | |
|---|---|---|
| positive regulation of cell death (GO:0010942) | 1.74 | 1.93E-02 |
| cell-cell signaling (GO:0007267) | 1.74 | 7.76E-03 |
| neurogenesis (GO:0022008) | 1.72 | 3.90E-06 |
| positive regulation of cell differentiation (GO:0045597) | 1.71 | 1.30E-03 |
| nervous system development (GO:0007399) | 1.71 | 1.30E-07 |
| negative regulation of transcription by RNA polymerase II (GO:0000122) | 1.7 | 6.67E-03 |
| regulation of cell differentiation (GO:0045595) | 1.7 | 2.69E-06 |
| cellular amide metabolic process (GO:0043603) | 1.69 | 4.81E-02 |
| negative regulation of transcription, DNA-templated (GO:0045892) | 1.62 | 3.26E-03 |
| negative regulation of nucleic acid-templated transcription (GO:1903507) | 1.62 | 3.59E-03 |
| negative regulation of RNA biosynthetic process (GO:1902679) | 1.62 | 3.65E-03 |
| regulation of developmental process (GO:0050793) | 1.61 | 4.58E-07 |
| negative regulation of cellular macromolecule biosynthetic process (GO:2000113) | 1.61 | 1.54E-03 |
| regulation of anatomical structure morphogenesis (GO:0022603) | 1.6 | 1.31E-02 |
| negative regulation of biosynthetic process (GO:0009890) | 1.6 | 9.57E-04 |
| regulation of multicellular organismal development (GO:2000026) | 1.6 | 1.75E-05 |
| chromosome organization (GO:0051276) | 1.6 | 3.29E-02 |
| negative regulation of macromolecule biosynthetic process (GO:0010558) | 1.6 | 1.66E-03 |
| negative regulation of RNA metabolic process (GO:0051253) | 1.59 | 3.26E-03 |
| negative regulation of cellular biosynthetic process (GO:0031327) | 1.59 | 1.47E-03 |
| negative regulation of nucleobase-containing compound metabolic process (GO:0045934) | 1.58 | 2.31E-03 |
| negative regulation of gene expression (GO:0010629) | 1.57 | 2.63E-04 |
| positive regulation of cellular component organization (GO:0051130) | 1.55 | 1.67E-02 |
| positive regulation of developmental process (GO:0051094) | 1.54 | 3.76E-03 |
| anatomical structure morphogenesis (GO:0009653) | 1.52 | 3.09E-04 |
| regulation of multicellular organismal process (GO:0051239) | 1.51 | 1.57E-06 |
| regulation of cellular component organization (GO:0051128) | 1.51 | 1.17E-04 |
| negative regulation of metabolic process (GO:0009892) | 1.5 | 1.43E-05 |
| negative regulation of macromolecule metabolic process (GO:0010605) | 1.5 | 5.54E-05 |
